# aDISCO: A clearing method to enable 3D microscopy of large archival paraffin-embedded human tissue blocks

**DOI:** 10.1101/2025.05.21.655358

**Authors:** Anna Maria Reuss, Dominik Groos, Martina Cerisoli, Lena Nordberg, Lukas Frick, Fabian F. Voigt, Nikita Vladimirov, Philipp Bethge, Regina Reimann, Fritjof Helmchen, Peter Rupprecht, Adriano Aguzzi

## Abstract

Human surgery and autopsy specimens are routinely stored as formalin-fixed paraffin-embedded (FFPE) tissue blocks. Diagnoses are based on microscopic examination of two-dimensional sections. Good clinical practice requires that samples be retained for decades, thus giving rise to vast archives of healthy and diseased tissues. While tissue clearing and whole-mount microscopy enable 3D analysis, FFPE human tissue blocks are often unsuitable for clearing and immunolabeling due to their large size, extensive cross-linking, and wax embedding. Here, we introduce ‘archival’ DISCO (aDISCO), a versatile and robust clearing method designed to overcome these challenges. aDISCO achieves complete clearing and immunolabeling of large samples stored for 15 years or more. We applied aDISCO to human brain, spinal cord, peripheral nerve, skin, muscle, heart, kidney, liver, spleen, colon, and lung, using a broad range of antibodies. To investigate the clinical usefulness of quantitative 3D histology by aDISCO, we performed a case study of focal cortical dysplasia, a neurodevelopmental disease causing epilepsy. Using deep-learning-based analysis, we found disrupted cortical layering and both focal and global neuronal density variations in FCD, features that are likely to be overlooked by conventional histology. In summary, aDISCO delivers large datasets suitable for deep-learning-based processing, enabling the detection of subtle and sparse pathologies in large archival human tissue specimens.

## Introduction

Histology is an ubiquitous tool for the diagnosis of diseases and for biological research (*1, 2*). Formalin-fixed, paraffin-embedded (FFPE) tissue preservation was pioneered by Ferdinand Blum in 1893 (*3*) and became widely adopted in the early 20^th^ century. FFPE tissue blocks are usually sliced into thin sections measuring about 5 µm (*4*). FFPE histology generates optically clear sections accessible to transmission light microscopy. However, such analyses are intrinsically two-dimensional and enable examination of only a small fraction of the tissue specimen. This, in turn, may prevent the detection of important pathologies and can only partially be obviated by the laborious scrutiny of numerous layered sections. These limitations impair the evaluation of spatially complex, convoluted structures such as blood vessels and nerve fibers.

3D histology has the potential to transcend these limitations. However, it requires transparency of thick tissue blocks, since the opacity inherent to most biological tissues (resulting from light scattering by complex internal structures such as lipids with different refractive indices (RIs) and light absorption by pigments) impedes microscopic examination (*5*). To overcome these physical limitations, lipids are removed, and RIs are matched to the immersion medium. While various tissue clearing methods were invented and have evolved in recent years, human tissue clearing presents unique challenges (*6*).

Since the dimensions of human organs exceed those of common model organisms, human tissue samples can be large. Typical FFPE human tissue blocks (∼2.5 cm x ∼2 cm x up to 1 cm) exceed the size of a mouse brain. Moreover, tissues rich in lipids and collagen such as brain white matter and liver are difficult to clear and stain effectively (*7–9*). Also, tissue fibrosis, which increases physiologically with age, makes sample preparation from aged adult patients particularly challenging (*10, 11*). Thus, processing human samples for 3D histology must achieve effective clearing without compromising tissue integrity and enable complete antibody diffusion without major staining gradients towards the center of the tissue specimen. Both issues are particularly challenging in archival FFPE tissues due to aldehyde cross-linking and epitope masking through formalin fixation, dehydration, and paraffin substitution (*12*). Finally, immunofluorescent staining of human samples entails additional issues such as autofluorescence from pigments like lipofuscin, melanin, or coagulated blood, which can disturb the desired staining signals (*6*).

For these reasons, current human tissue clearing methods with successful immunolabeling are limited to either small (< 2 mm thick) FFPE samples or freshly processed samples from few organs, primarily from the brain (*13–28*). A robust 3D histology pipeline for effective clearing and immunolabeling of large, long-term-stored FFPE human tissue specimens that is applicable across various organs and coupled with a validated set of antibodies remains a major unmet need.

Here, we introduce ‘archival DISCO’ (aDISCO), a versatile and robust clearing method designed to achieve effective clearing and antibody staining in whole-mount FFPE human tissue blocks from diagnostic archives, thereby making vast collections of human tissue samples available for deeper investigation (Fig. 1). We demonstrate the suitability of aDISCO across different human organs and its compatibility with a broad set of antibodies, obtaining whole-mount 3D images at various magnifications using light-sheet fluorescence microscopy (mesoSPIM versions (V)5 and V6) (*29, 30*). In a use-case study based on a relatively rare pathological condition (focal cortical dysplasia, FCD), we demonstrate that 3D histology with aDISCO generates high-quality data that can be processed with automated segmentation and object detection methods based on deep-learning. Our results showcase the potential of quantitative 3D histology with aDISCO to detect subtle and focal pathologies in large archival human tissue specimens.

**Fig. 1.**
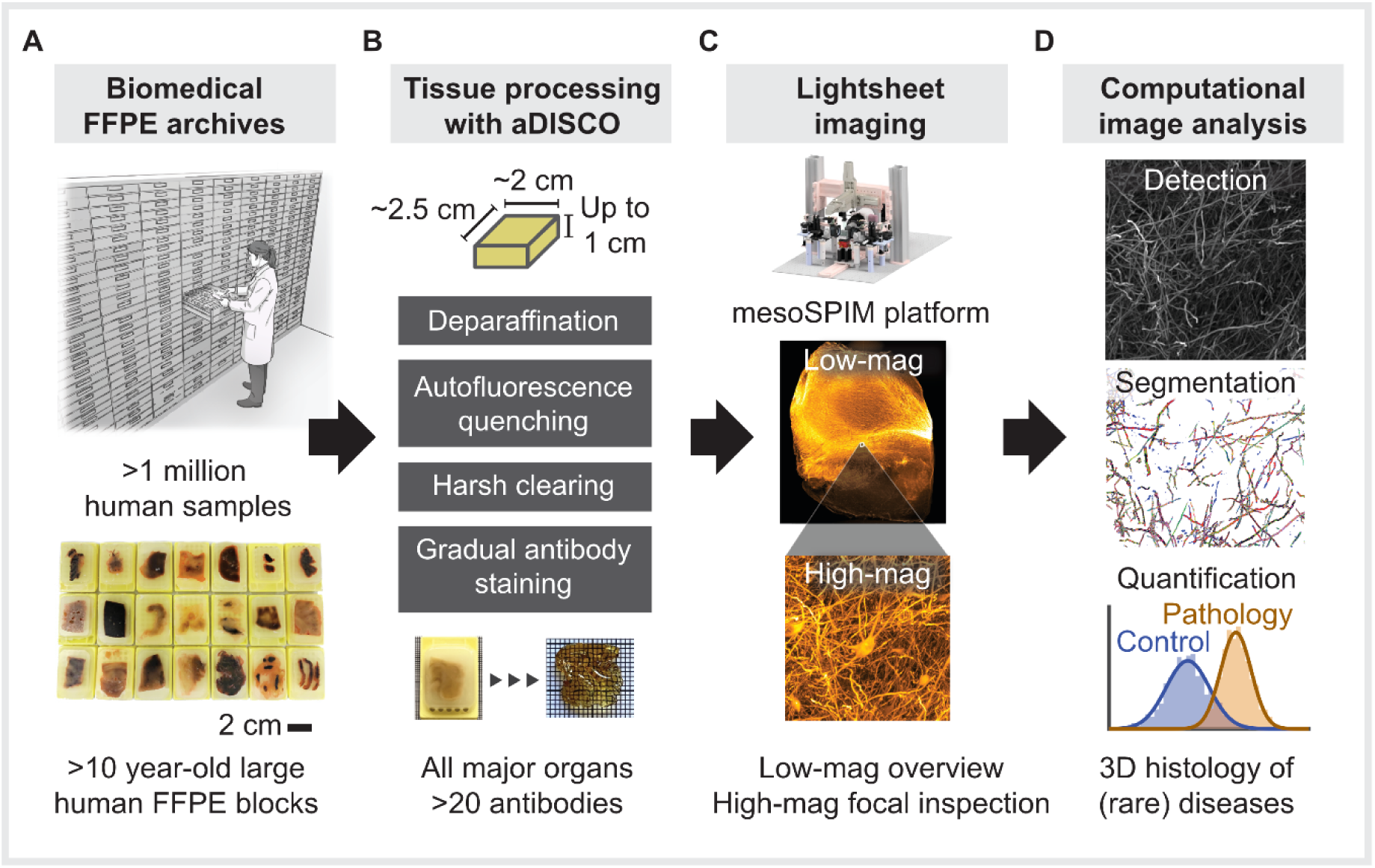
3D histology workflow for entire archival human FFPE tissue blocks using aDISCO. (**A**) Biomedical FFPE archives contain more than a million human samples from all major organs and various pathological backgrounds, often stored for decades. (**B**) aDISCO is designed for processing large human FFPE tissue blocks, and was tested on all major organs and with more than 20 antibodies. (**C**) Light-sheet imaging with the mesoSPIM enables low-magnification (mag) overviews and high-mag focal inspection of cleared and immunolabeled human tissue blocks at (sub-)cellular resolution. (**D**) aDISCO enables quantitative 3D histology of healthy human tissues and (rare) pathologies from FFPE samples using deep learning-based detection and segmentation.

## Results

### Development of aDISCO

We developed a clearing protocol that enables the processing of long-term-stored (samples used in this study stored for 11-15 years) standard FFPE tissue blocks from various human organs, which are customarily stored in diagnostic archives for surgical pathology, autoptic pathology, and at institutes of human anatomy (Fig. 1, Fig. 2A). The method presented here is based on three main innovations: (1) harsh clearing with a combination of tetrahydrofuran (THF) and dichloromethane (DCM), (2) strong autofluorescence suppression using acidic cupric sulfate in ammonium acetate, and (3) gradual immunolabeling with increasing antibody concentrations.

**Fig. 2.**
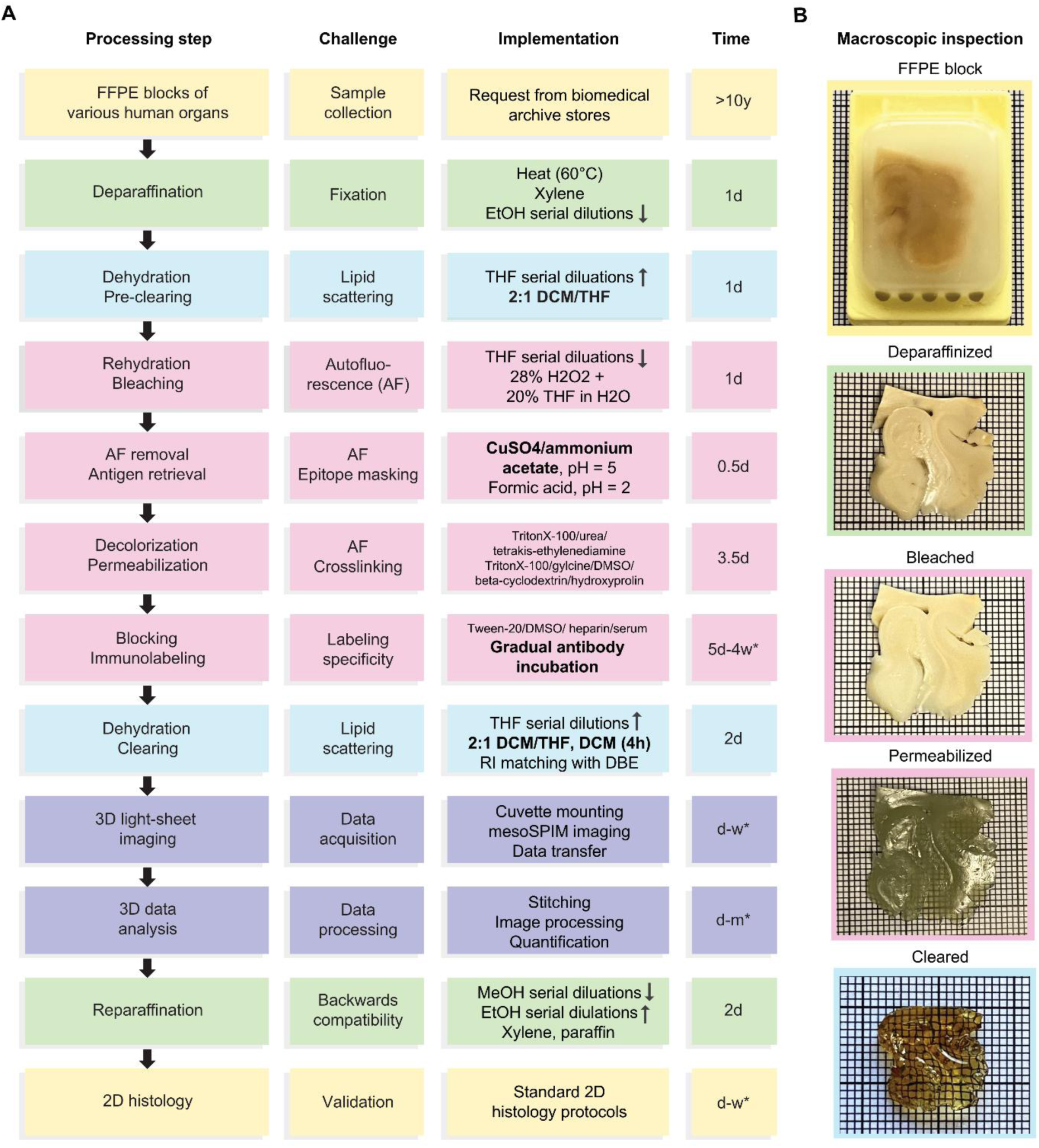
Overview of the aDISCO protocol for clearing and immunolabeling of archival human tissue blocks. (**A**) The aDISCO pipeline (first column), with the associated challenges of processing large archival human tissue blocks (second column), the implementations to overcome these challenges (third column), and the typical times required for the steps (fourth column). FFPE human tissue blocks from biomedical archive stores are processed using harsh deparaffination, pre-clearing, and clearing conditions with a novel mixture of dichloromethane (DCM) and tetrahydrofuran (THF) to reach optical transparency. A high concentration of hydrogen peroxide (H_2_O_2_) and cupric sulfate (CuSO_4_) in ammonium acetate remove autofluorescence (AF). Antigen (Ag) retrieval with formic acid helps unmask epitopes after overfixation. Together with pre-clearing, decolorization, and intense permeabilization, gradual antibody incubation leads to effective antibody diffusion throughout the large and dense tissue blocks. After 3D light-sheet imaging (mesoSPIM V5 and V6 (*30*)) and 3D data analysis, cleared samples can be reparaffinized and reused for conventional 2D histology. DBE: dibenzylether, DMSO: dimethylsulfoxide, EtOH: ethanol, H2O: water, RI: refractive index, d: days, m: months, w: weeks. (**B**) Quality control by macroscopic inspection of an archival human brain sample after serial procedures as described. Small squares: 1 mm.

Standard FFPE human tissue blocks are relatively large, with the exact sizes depending on sampling and prior cutting. In addition, various tissues including skin, liver, and white matter-rich regions of the brain are very dense (*8, 9, 31*). To address the high density and extensive cross-linking in large FFPE human tissue blocks, particularly in lipid and collagen-rich organs, we developed a harsh clearing strategy utilizing a combination of the polar and organic solvents THF and DCM (Fig. 2A). These chemicals are used repeatedly to achieve stepwise delipidation. In addition, the pre-clearing removes the barriers to molecular flux inf dense tissues (*5*), which facilitates antibody penetration for immunolabeling.

Traditional bleaching agents like hydrogen peroxide, frequently used in other clearing protocols (*10, 15*), often prove insufficient in suppressing the strong autofluorescence from coagulated blood, lipofuscin, melanin, and other pigments (*6*) inherent to human archival tissues (Fig. S1A). This persistent autofluorescence is problematic for automated segmentation during 3D data analysis, because brightly autofluorescent artifacts may be falsely interpreted as positive staining signals. To overcome this limitation, aDISCO incorporates a robust autofluorescence suppression step using acidic cupric sulfate in ammonium acetate, whose effectiveness had been demonstrated on 2D sections already in the 1990s (*32*). Crucially, we found that its application after immunolabeling suppressed the desired staining signal (Fig. S1B). Therefore, we applied it before initiating the staining procedure (Fig. 2A), thus preserving the specific staining signal while removing autofluorescence background (Fig. S1B).

One of the most difficult challenges in 3D histology of large, dense archival human tissues is to achieve full antibody penetration and staining throughout the entire blocks. Therefore, immunolabeling has only been achieved in either freshly processed human samples (*13, 17, 18, 20–28*) or small FFPE human tissue specimens with limited imaging depth (*14, 19, 28*) (Table 1).

**Table 1.** Comparison of human tissue clearing methods.

|  | iDISCO<br>(13) | DIPCO<br>(14) | MASH<br>(15, 16) | SHANEL | aDISCO | OPTI<br>Clear<br>(19) | FAST<br>Clear<br>(20, 21) | CLARITY<br>(22, 23) | SWITCH<br>(24)<br>SHIELD<br>(25) | HIF<br>(26) | ELAST<br>(27) | CUBIC<br>(28) |
| --- | --- | --- | --- | --- | --- | --- | --- | --- | --- | --- | --- | --- |
| **FFPE** | no | yes | yes | no | **yes** | yes | no | no | no | yes | no | yes |
| **Long-term**<br>**formalin-**<br>**fixed** | no | no | yes | no | **yes** | yes | no | yes | yes | no | yes | no |
| **Imaging**<br>**depth** | < 1 mm | 1.6 mm | 3 mm<br>(no Abs) | 1 cm | **6.5 mm** | 546 μm | < 509 μm | 2.6 mm | 100 μm | 900 μm | 5 mm | 2.1 mm<br>(FFPE<br>no Abs),<br>400 μm<br>(fresh) |
| **Full**<br>**antibody**<br>**penetration** | yes | yes | N/A<br>(no Abs) | no | **yes** | yes | yes | no | yes | yes | no | NA<br>(FFPE<br>no Abs),<br>yes<br>(fresh) |
| **No. of**<br>**human**<br>**organs** | 1 | 6 | 2 | 7 | **11** | 1 | 2 | 1 | 1 | 1 | 1 | 2 |
| **No. of**<br>**primary**<br>**antibodies** | 2 | 5 | 0 | 10 | **22** | 4 | 3 | 9 | 10 | 3 | 13 | 0<br>(FFPE),<br>2 (fresh) |

FFPE: Formalin-fixed paraffin-embedded, Ab: antibody, No.: number, N/A: not applicable.

Previous clearing protocols used a single antibody concentration, which was often applied at high concentration to reach bright staining signals (*13, 14, 17*). However, we observed that in large samples this procedure typically results in a bright rim and insufficient staining inside the tissue, with a visible signal gradient or even a dark center. To prevent accumulation of antibodies at the tissue rim, we employed an approach starting with low antibody concentrations, which we gradually increased by adding more and more antibodies to the staining solution (Fig. 2A). In addition, we introduced a series of consecutive permeabilization steps before and during the immunolabeling procedure by combining various reagents which have been previously shown to facilitate antibody diffusion (*10, 33–35*) (Fig. 2A). Furthermore, epitopes are often masked in FFPE tissue by overfixation and paraffin embedding, preventing the antibodies from binding to their respective antigens (*12*). For antibodies yielding weak staining signals, we performed antigen retrieval with formic acid (Fig. 2A).

The combination of these innovations enabled reliable clearing and immunolabeling of entire FFPE tissue blocks from various human organs and was compatible with a substantial set of commonly used antibodies to characterize these organs.

### Validation of 3D histology with aDISCO-processed whole-mount archival human tissue blocks

Validation of 3D findings by 2D histology requires clearing of archival tissue to be compatible with subsequent re-embedding in paraffin, conventional histological processing, and staining (Fig. 2A). We developed a reparaffination protocol which gently restores the original FFPE tissue block integrity and results in 2D sections with similar staining quality as the original non-cleared FFPE tissue sections (Fig. S1C).

Tissue clearing procedures are laborious and can take several weeks depending on sample size. It is therefore helpful to assess their performance at intermediate stages. We performed such quality testing on archival human brain samples during aDISCO processing (Fig. 2B). After deparaffination, archival human tissue is usually of brownish color, which becomes whitish after successful bleaching and greenish after cupric-sulfate treatment. After strong permeabilization and decolorization, less compact tissues such as the cortical gray matter of the brain already start becoming transparent, whereas the white matter remains opaque. Final clearing and refractive index (RI) matching render the entire sample fully transparent.

### aDISCO enables high-quality clearing and immunolabeling throughout large, dense human tissue blocks

To assess the performance of the aDISCO protocol, we processed an archival human midbrain sample with a high content of lipid-rich white matter, stored as FFPE for >10 years. We stained the sample with anti-tyrosine hydroxylase (TH) antibodies which visualize dopaminergic neurons (Fig. 3A, Movie S1). In a 3D rendering of the whole-mount imaged sample using light-sheet microscopy, the architecture of the human midbrain, including the substantia nigra, can be identified (Fig. 3A). Region-of-interest (ROI) light-sheet imaging of the substantia nigra with higher magnification (Fig. 3B and C) highlights branching neurites, including fine dendrites.

**Fig. 3.**
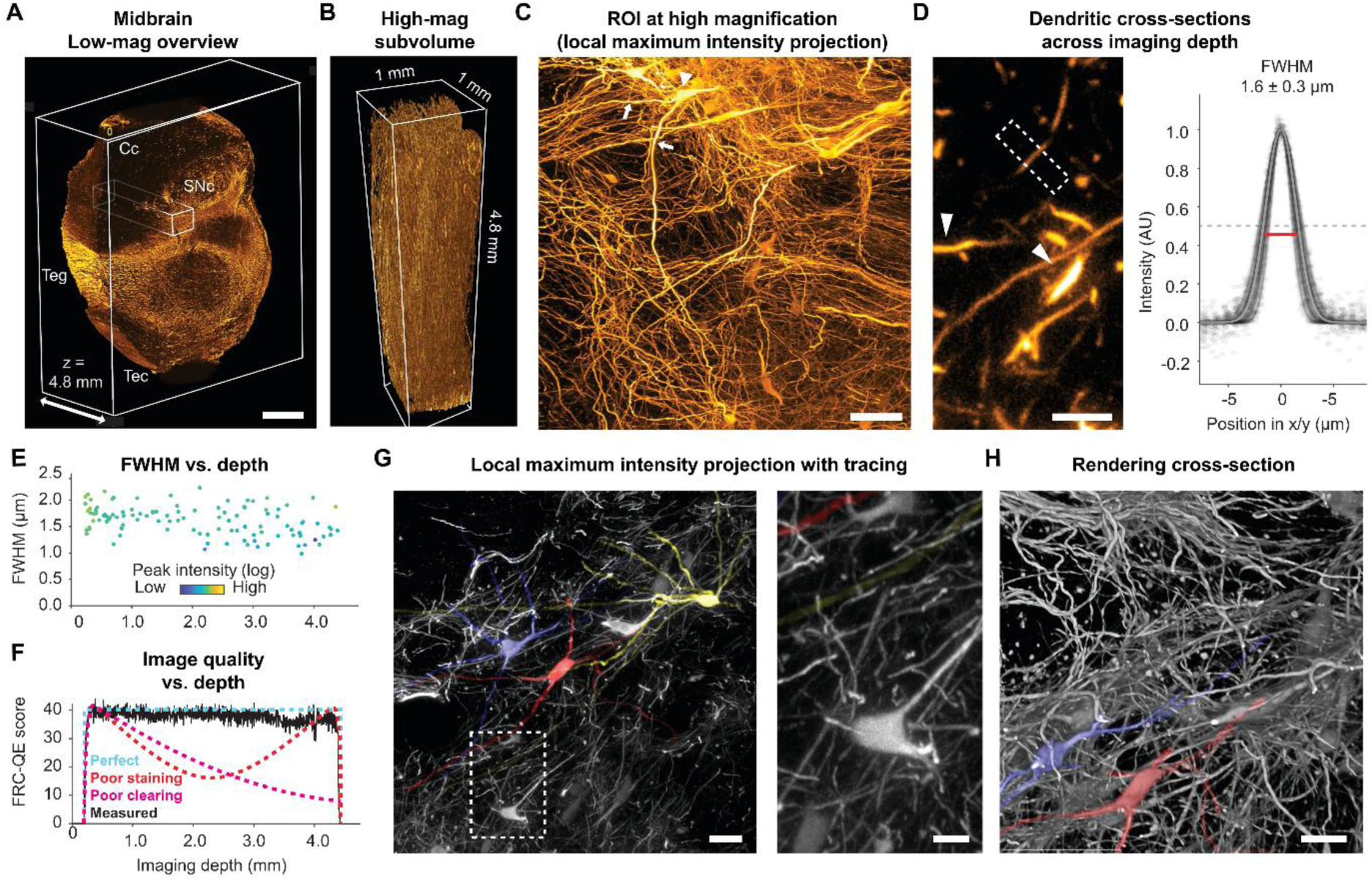
Large-scale screening and high-resolution analysis of archival human midbrain. (**A**) 3D rendering of archival human midbrain, stained for dopaminergic neurons (anti-tyrosine hydroxylase antibody), and imaged at low optical magnification (mag; 1.25x) using mesoSPIM V5. The rendering shows the crus cerebri (Cc), tegmentum (Teg), tectum (Tec), and substantia nigra pars compacta (SNc). Scale bar: 2 mm. (**B**) Subvolume of the SNc imaged at high mag (12.6x) using mesoSPIM V5. (**C**) Local maximum intensity projection of the high-mag subvolume highlights large dopaminergic neurons with somata (arrowhead) and long neurites (arrows), branching into a dense fiber network. Scale bar: 50 µm. (**D**) Left: Thin dendrites (dashed white rectangle) are selected for cross-sectional profiling as a readout of lateral resolution across sample depth. In contrast, bright and thick dendrites (arrowheads) are avoided for analysis. Right: Dendritic cross-sectional intensity profiles with Gaussian fits in arbitrary units (AU; n = 108 across all depths). The full width at half maximum (FWHM, median ± standard deviation) was obtained from Gaussian fits to individual profiles. **(E**)The lateral FWHM shows no depth-dependent increase. Peak intensities of dendritic profiles are color-coded (logarithmically spaced in arbitrary units. (**F**) Fourier ring correlation quality estimate (FRC-QE) across imaging depth. In an ideal case, FRC-QE would be constant across imaging depth (light blue). For poorly stained samples, FRC-QE would be high at the sample edges and low in the center (red). For poorly cleared samples, FRC-QE would decrease with imaging depth (magenta). The measured FRC-QE is stable across imaging depth (black). (**G**) Local maximum intensity projections with neuronal tracing across 30 µm. The clearing and staining quality allow for tracing of the main neuronal processes in the sample center (imaging depth approx. 2 mm), as shown here for three example neurons with distinct color coding. The inset highlights small neuronal processes. (**H**) 3D rendering of traced neuronal processes. See Movie S2.

Optical inhomogeneities lead to light scattering and result in a reduction of the effective imaging resolution. The nominal lateral resolution of our imaging method is 1.7 µm. To demonstrate effective clearing and immunolabeling throughout sample depth, small and ideally subdiffractive objects can be inspected. To this end, we analyzed the cross-sectional profiles of small neuronal dendrites across depth (Fig. 3D), with no obvious increase of dendritic cross-sectional width with depth, indicating preserved lateral resolution throughout the full thickness of the cleared sample (Fig. 3E). The variability of measured dendritic width can be explained by varying peak intensities of the inspected dendrites, indicating that thicker dendrites slightly broadened the profiles due to their thickness or due to saturation effects. A complementary analysis based on Fourier ring correlation analysis (*36*) confirmed that image quality was relatively stable across imaging depth (Fig. 3F). To demonstrate that this high image quality enables the analysis of subcellular structures, we used the neuronal tracing framework webKnossos (*37*) to track the main processes of selected neurons within the center of the high-magnification subvolume (Fig. 3G and H, Movie S2). Together, these analyses indicate that aDISCO enables imaging at high, even subcellular resolution across the entire depth of lipid-rich archival human tissue blocks.

### 3D histology of different human central nervous system (CNS) regions

Next, we tested the applicability of aDISCO across a broad spectrum of human archival CNS samples (Fig. 4, Movies S3 and S4). We used various antibody stains and light-sheet microscopy to visualize NeuN-positive neurons or specifically parvalbumin-positive interneurons in the cortex, IBA1-positive microglia and podocalyxin-positive blood vessels in the hippocampus, tryptophane hydroxylase (TPH)2-positive serotonergic neurons in the brain stem, neurofilament-positive neurons, including Purkinje cells, and nerve fibers in the cerebellum, and GFAP-positive astrocytes and SMA-positive arterial vessels in the spinal cord.

**Fig. 4.**
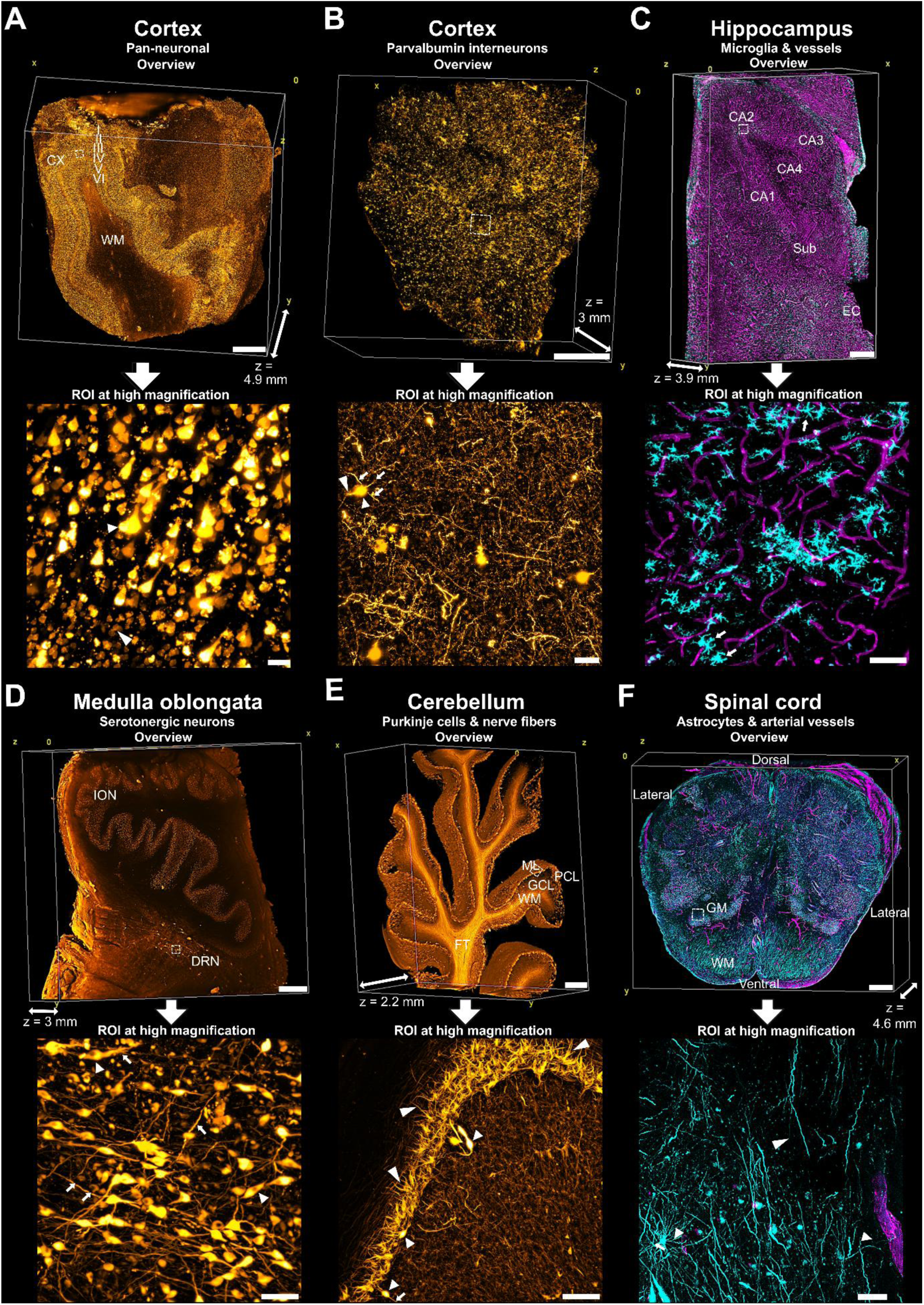
3D histology of different human CNS regions. Each panel presents a low-magnification 3D rendering of a whole-mount sample (top; 0.8x–2x) and a high-magnification local maximum intensity projection of a region-of-interest (ROI; bottom; 12.6x), imaged using mesoSPIM V5. The ROIs are indicated in the rendering by dashed white boxes. (**A**) Archival human brain cortex (CX) sample, stained for neurons (anti-NeuN antibody), revealing cortical layers I-VI and the white matter (WM). Scale bar: 2 mm. The ROI demonstrates neuronal cytoarchitecture of granular cells (large arrowhead) and large pyramidal cells (small arrowhead) in layer III. Scale bar: 50 µm. (**B**) Parvalbumin-positive interneurons in archival human brain cortex. Scale bar: 1 mm. The ROI shows neuronal somata (small arrowhead) with axons (large arrowhead) and dendrites (arrows). Scale bar: 100 µm. (**C**) IBA1-positive microglia (cyan) and podocalyxin-positive blood vessels (magenta) in aged archival human hippocampus across distinct anatomical subregions (cornu ammonis (CA)4, CA3, CA2, CA1, subiculum (Sub), and entorhinal cortex (EC)). Scale bar: 1 mm. The ROI reveals microglia morphology with their short processes (arrows) close or attached to blood vessels. Scale bar: 50 µm. (**D**) Tryptophan hydroxylase (TPH)2-positive serotonergic neurons in archival human medulla oblongata. Distinct lower brainstem nuclei such as the inferior olivary nucleus (ION) and the dorsal raphe nucleus (DRN) are visible. Scale bar: 1 mm. The ROI in the DRN displays serotonergic neurons with nuclei (arrowheads) and widely ramified neurites (arrows). Scale bar: 50 µm. (**E**) Archival human cerebellum, stained with anti-neurofilament antibody to visualize cerebellar anatomy with fiber tracts (FT) of the WM and distinct cerebellar cortical layers, consisting of the molecular layer (ML), Purkinje cell layer (PCL), and granular cell layer (GCL). Scale bar: 1 mm. The ROI displays Purkinje cells with somata (small arrowheads), arborized dendrites (large arrowheads), and axons (arrow). Scale bar: 50 µm. (**F**) GFAP-positive astrocytes (cyan) and SMA-positive arterial vessels (magenta) in archival human spinal cord. The characteristic butterfly-like shape of the grey matter (GM) can be distinguished from the white matter (WM) due to distinct astrocytic staining patterns. Scale bar: 1 mm. The ROI displays stellate astrocytes (arrow) and long glial processes (large arrowhead) besides arterial vessels (small arrowheads). Scale bar: 50 µm.

To assess the clearing and immunolabeling quality in the sample center, we performed orthogonal reslicing of an anti-NeuN-labeled human cortex sample that exhibits an evenly distributed staining pattern across the entire cortex and is therefore suitable for this type of analysis (Fig. S2A). In all three image projections, staining signalsin the cortex were preserved throughout the reslices. In addition, both cortex and lipid-rich white matter did not exhibit obvious signs of blurrying, indicating effective clearing of the entire sample.. Next, we calculated the mean fluorescence intensity (MFI) projection along the z-direction of the entire sample. In the x- and y-directions, MFI was similar between superficial and deep areas of similar composition, with inhomogeneities reflecting region-specific cellular distributions rather than staining gradients (Fig. S2B; see Fig. S3 for MFI projections of the other CNS samples). In addition, we computed the z-profile of the mean fluorescence intensity averaged across x and y. The z-profile remained stable throughout the entire range in the vertical (z) direction, even in the center of the sample (Fig. S2C). In general, imaging depth was not limited by antibody penetration or clearing quality, but by the size of the respective tissue blocks (Fig. 3 and 4), which was 8.7 ± 2.8 mm (mean ± s.d. across n = 7 CNS samples) x 7.5 ± 1.9 mm, and 3.7 ± 1.0 mm thick after tissue shrinkage of about 30%, following organic solvent-based tissue clearing (*38*). Accordingly, the maximum imaging depth was 4.9 mm, corresponding to the entire sample thickness of the anti-NeuN-stained archival human cortex sample after clearing and accompanying tissue shrinkage (Fig. 4A).

To further make use of the high optical resolution enabled by aDISCO, we imaged ROIs across all processed CNS samples at higher magnification (insets in Fig. 4). This allowed us to identify individual somata and neurites of several neuronal cell types (Fig. 4A, B, D, and E), the interactions of microglia feet with finely branched blood vessels (Fig. 4C), and the processes of stellate astrocytes in the spinal cord (Fig. 4F).

In summary, we found that aDISCO achieves effective clearing and immunolabeling of large archival human samples across the entire CNS spectrum, while enabling subcellular resolution of fine structures even deep within each tissue block. Since the staining quality did not degrade after up to 15 years of storage (anti-NeuN-labeled cortex sample), aDISCO seems suitable for the investigation of historical samples.

### 3D histology of human organs beyond the CNS

In principle, aDISCO should be applicable to human organs other than the CNS, but adaptations may be needed because of differences in tissue composition. We therefore processed whole-mount FFPE human tissue blocks from disparate organs, stained them with a panel of antibodies, and imaged them by light-sheet microscopy (Fig. 5,Movies S5 and S6). With aDISCO, we visualized myelinated fibers in peripheral nerve, myelinated and unmyelinated nerve fibers in the skin, skeletal muscle fibers together with their vascularization, cardiac myocytes and arteries in the heart, glomeruli and tubules in the kidney, sinusoids in the spleen, bile ducts in the liver, bronchioles and alveoli in the lung, and the mucosa with crypts in the colon.

**Fig. 5.**
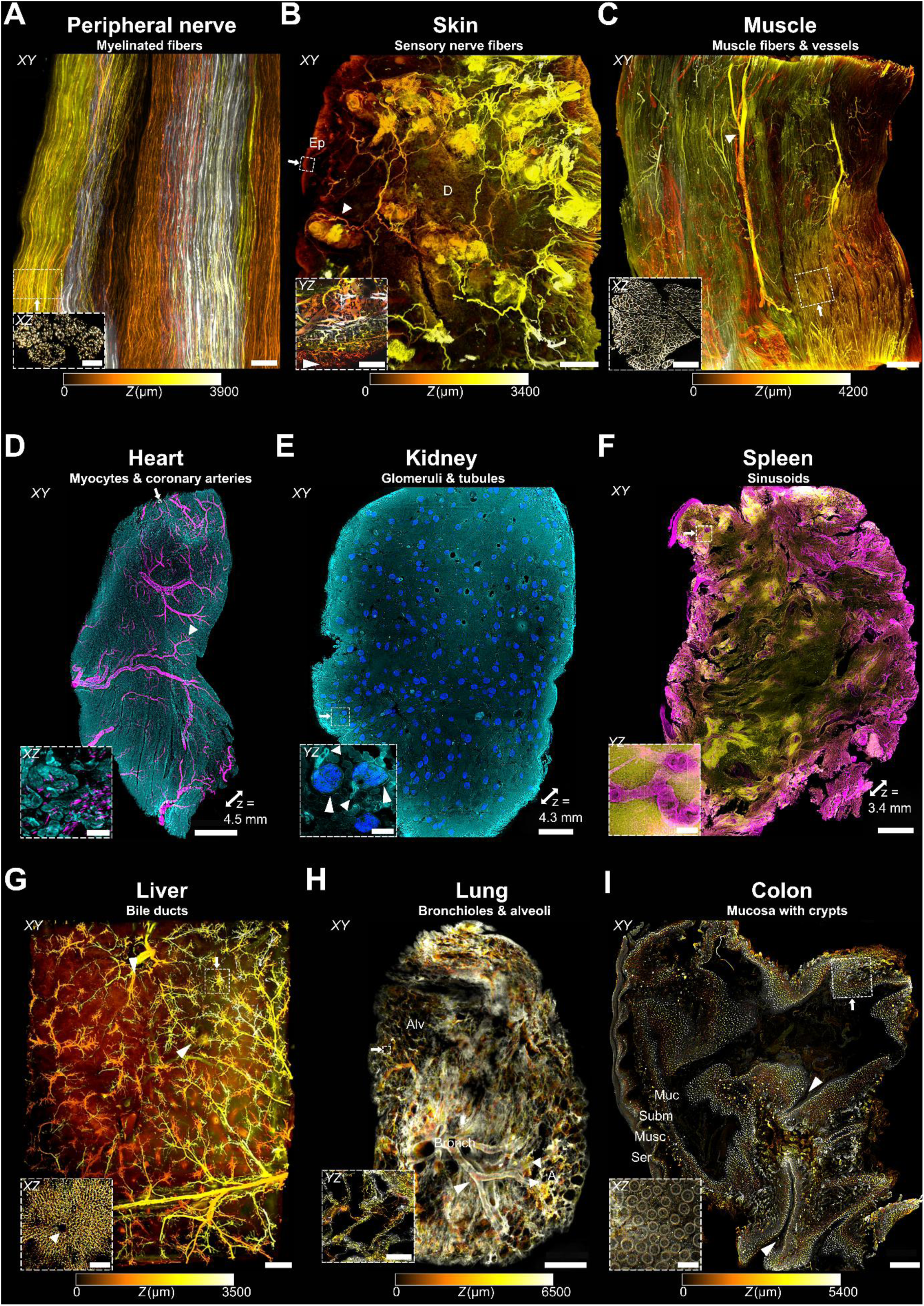
3D histology of human organs beyond the CNS. Each panel displays an xy-projection for sample overview, imaged at 0.63x-12.6x magnification using mesoSPIM V5. Panels A-C and G-I are depth color-encoded to show the three dimensions of each sample. Panels D-F are local maximum-intensity projections with indicated z-thickness of each sample to illustrate multicolor antibody staining. The insets in each panel (bottom left) display orthogonal sections each of a region-of-interest (ROI) from the same sample imaged at higher magnification (4x-20x using mesoSPIM V5 and V6) to demonstrate high axial resolution in z. The ROIs are indicated in the depth color-encoded overview by dashed white boxes and the direction of the ROI viewport is illustrated by the arrow next to the white box. (**A**) MBP-positive myelinated fibers in archival human peripheral nerve. The longitudinal view displays the cotton wool-like pattern of thin nerve fibers. Scale bar: 500 µm. The inset highlights the nerve fascicles composed of individual myelinated fibers in the transversal view. Scale bar: 100 µm. (**B**) PGP9.5-positive sensory nerve fibers in archival human skin. The longitudinal view displays branching nerve fibers in the epidermis (Ep) and dermis (D), forming Pacinian corpuscles (small arrowhead). Scale bar: 1 mm. The inset highlights the innervation pattern of the epidermis by thin sensory nerve terminals (large arrowhead). Scale bar: 100 µm. (**C**) Laminin-positive muscle fibers and blood vessels in archival human skeletal muscle with prominent vascularization (arrowhead in the longitudinal view). Scale bar: 1 mm. The inset highlights the muscle fascicles composed of individual muscle fibers in the transversal view. Scale bar: 200 µm. (**D**) N-cadherin-positive myocytes (cyan) and SMA-positive coronary arteries (magenta) in archival human heart. The longitudinal view highlights the branching of coronary arteries (arrowhead) in the myocardium. Scale bar: 2 mm. The inset displays a ROI xz-projection of smaller arterioles between myocytes. Scale bar: 30 µm. (**E**) Nephrin-positive glomeruli and E-cadherin-positive tubules in archival human kidney. Scale bar: 1 mm. The inset displays a ROI yz-projection with detailed morphology of glomeruli with Bowman capsules (large arrowheads) and outgoing proximal tubules (small arrowheads). Scale bar: 150 µm. (**F**) SMA-positive sinusoids in CD45-positive lymphatic tissue of archival human spleen. Scale bar: 1 mm. Inset scale bar: 100 µm. (**G**) CK19-positive bile ducts in archival human liver, displaying the branching pattern (large arrowheads) of hepatic bile ducts in the longitudinal view. Scale bar: 1 mm. The inset shows a hepatic lobule with central vein (small arrowhead) surrounded by bile ducts in the transversal view. Scale bar: 200 µm. (**H**) CK7-positive airways in archival human lung. In the longitudinal view, the bronchiolar tree with bronchioles (Bronch), branching (large arrowhead) and segueing (small arrowheads) into peripheral alveoli (Alv) is visible. Scale bar: 2 mm. The inset displays a ROI yz-projection of alveoli. Scale bar: 100 µm. (**I**) Collagen-IV-positive archival human colon sample. The longitudinal view highlights deeply caved mucosal crypts (arrowheads) within the colon wall composed of mucosa (Muc), submucosa (Subm), muscularis (Musc), and serosa (Ser). Scale bar: 1 mm. The inset shows the arrangement of individual crypts within the colon wall structure, from the inner mucosa to the outer serosa (double arrow) in the transversal view. Scale bar: 300 µm.

These examples demonstrate that aDISCO achieves effective clearing and immunolabeling of cellular structures across a broad variety of human organs. This finding was supported by MFI projections, showing robust staining signals in each tissue block (Fig. S4). This was also true for very dense tissues, such as liver or skin. Again, imaging depth was not limited by antibody penetration or clearing quality, but by the size of the respective tissue blocks, which was 10.7 ± 4.7 mm (mean ± s.d. across n = 9 samples) x 7.6 ± 3.2 mm, and 4.3 ± 1.5 mm thick after tissue shrinkage of about 30%, following organic solvent-based tissue clearing (*38*). Accordingly, the maximum imaging depth was 6.5 mm, corresponding to the entire sample thickness of the anti-CK7-stained archival human lung sample after clearing and accompanying tissue shrinkage (Fig. 5H).

The clearing and staining quality enabled the resolution of thin structures (insets in Fig. 5), including the branches of sensory nerve terminals in the epidermis of the skin (inset in Fig. 5B, Movie S5) and the Bowman capsules and outgoing proximal tubules of glomeruli in the kidney (inset in Fig. 5E, Movie S6).

In conclusion, our findings highlight the versatility of aDISCO for 3D histology of whole-mount FFPE human tissue blocks across multiple organs and its compatibility with a broad range of antibodies for detailed organ characterization.

### 3D cortical profiling of neuronal density in archival human brain

Since large cleared samples can be imaged volumetrically using 3D microscopy, aDISCO may enable the three-dimensional quantitative morphometry of tissues such as human cortex with analytical precision that is unattainable by conventional histology. The human cortex exhibits a characteristic architecture consisting of six neuronal layers (*39*) and is essential for performing complex neuronal computations, which are the basis of cognition and behavior. Precise quantification of cortical neurons and their distribution across distinct layers may help advance the understanding of the relationships between functional integrity and cellular organization in the human cortex. This knowledge may be relevant to many fields of neuroscience, from refining models of neuronal computation to elucidating structural alterations during development, aging, and in pathological states.

The characterization of human cortical layers (*40*) relies primarily on 4-5 µm thin 2D sections (*41, 42*). In contrast, comprehensive 3D examination of large human cortical specimens has been limited to counting total cell numbers using small molecular dyes, which penetrate thick tissues more easily but do not distinguish neurons from other cell types (*16, 17*). In our study, specific immunolabeling of neurons with anti-NeuN antibody allowed us to precisely assess neuronal density in large archival human cortex specimens, enabling detailed three-dimensional profiling of the neuronal density across distinct cortical layers (Fig. 6).

**Fig. 6.**
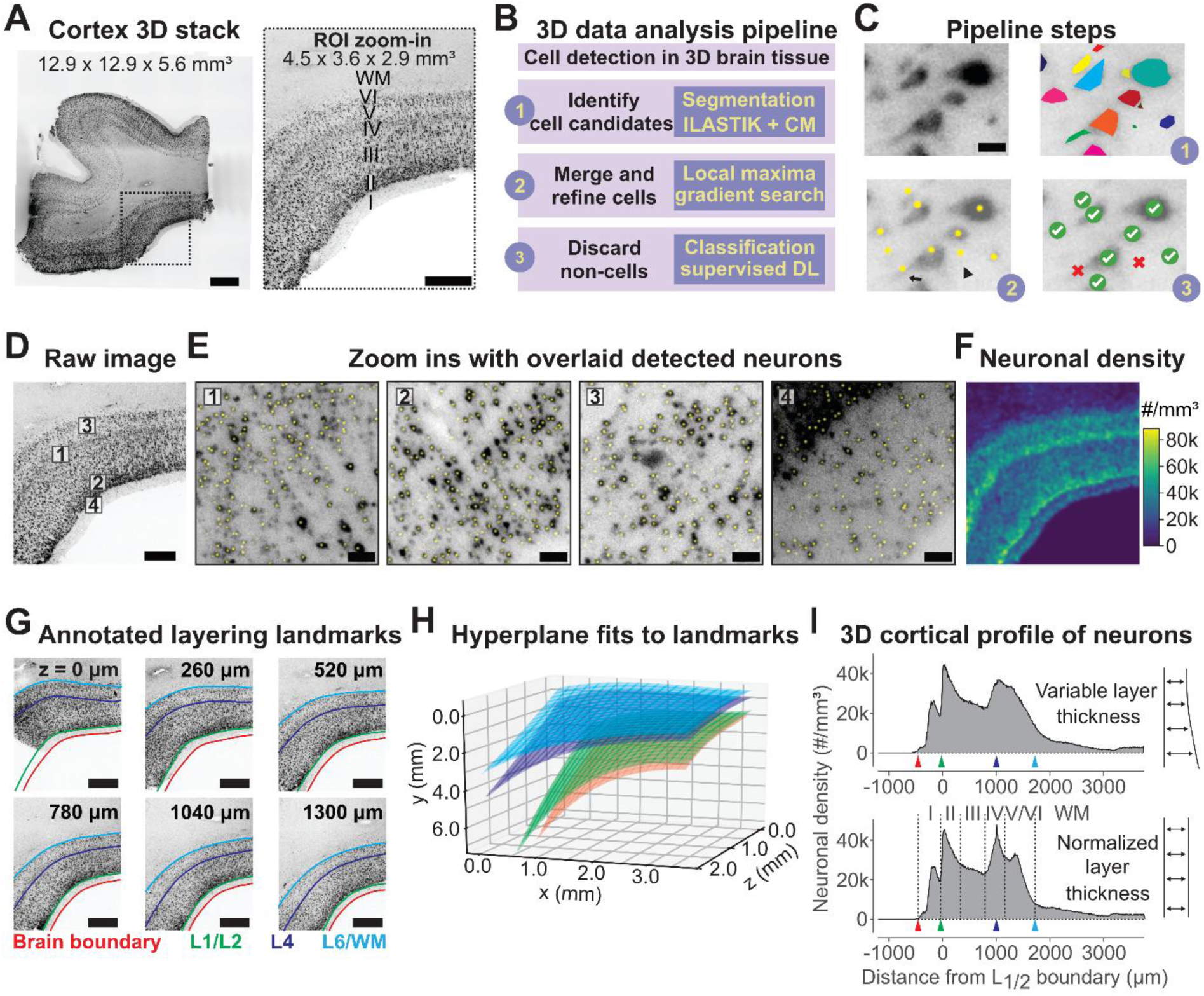
Neuronal density profiling in archival human brain cortex. (**A**) Left: Stitched and 4x-downsampled 3D stack of whole-mount FFPE human cortex specimen, stained for neurons (anti-NeuN antibody) and imaged at 4x magnification using mesoSPIM V5. Scale bar: 2 mm. Right: Digital zoom into a region-of-interest (ROI) to highlight cortical layers I-VI and the white matter (WM). Scale bar: 1 mm. (**B**) 3D data analysis workflow for neuronal cell detection in three steps. CM: ClearMap, DL: deep-learning. (**C**) Ilustration of the pipeline steps explained in (B). Upper left: Raw fluorescence image. Scale bar: 20 µm. Upper right: Segmented neuronal cell candidates, visualized by different colors. Lower left: Detection of cortical neurons based on local maxima (yellow dots) of segmented neuronal cell candidates. Note false positive detection of axons separated from somata (arrow) and of noise (arrowhead). Lower right: Falsely detected cell candidates are discarded (red crosses) via supervised DL, while correctly detected neurons are kept (green check marks). (**D**) Raw fluorescence image of cortical layers. Scale bar: 1 mm. (**E**) Digital zoom-in micrographs of regions numbered in (D) with overlaid detected neurons (yellow dots). Scale bars: 50 µm. (**F**) Spatial mapping of neuronal density. #: neuron count. (**G**) 3D annotation of visually identified layering landmarks for defining distinct layers (L), indicated by the different colors and exemplarily shown along the z-axis. Scale bars: 1 mm. (**H**) Generation of polynomial hyperplanes fitted to the respective layering landmarks. (**I**) Neuronal density per mm^3^ across all cortical layers, defined by the landmark hyperplanes in (H) as indicated by the colored arrows and plotted according to the distance from the boundary between layers 1 and 2 (L1/2). 3D cortical profile before (upper histogram) and after normalization to variable layer thickness (lower histogram).

We devised a 3D data analysis pipeline for efficient and accurate neuron counting in large datasets (Fig. 6B and C). We used ILASTIK (*43*) and ClearMap (*44*) for segmentation and detection of neuronal cell candidates, which were subsequently refined by merging falsely separated parts of neurons. Neuronal candidates were then classified as cells vs. non-cells, using a supervised deep convolutional neural network (DCNN) that used three orthogonal virtual slices of the local environment of each candidate to predict its class (cross-validated accuracy 92.2%; Fig. S5).

This approach enabled robust detection of neurons and subsequent determination of neuronal density in human cortex (Fig. 6D-F, Movie S7). To define cortical layers, we used visual guidance to annotate landmarks that indicate specific layers or boundaries between layers and fitted them to polynomial hyperplanes. This approach enabled precise determination of neuronal densities across distinct cortical layers (Fig. 6G-H). To compensate for variable cortical thickness within a 3D sample, we normalized layer thickness by the average distance between landmarks. As a result, we obtained a layer-dependent neuronal density profile which can distinguish distinct cortical layers, except layers 5 and 6 where the transition is also morphologically less well-defined (Fig. 6I).

In summary, our analyses show that large archival human samples processed via aDISCO provide clearing and antibody staining quality that is amenable to segmentation and object detection. Our 3D data analysis pipeline allows for accurate detection of neurons and delineation of neuronal densities across distinct cortical layers in three spatial dimensions. By establishing a reference profile of neuronal cell distribution in normal human cortex, we provide a basis for assessing cortical abnormalities under pathological conditions.

### 3D histological investigation of focal cortical dysplasia (FCD)

To test a use case for these new tools, we investigated the cortical architecture in FCD, a neurodevelopmental malformation disorder causing pharmaco-resistant epilepsy in children and young adults (*45*). Its prevalence within the general population is < 1% (*46*), but ranges between 5% to 25% among patients suffering from focal epilepsy (*47*). FCD is characterized by distortions in cortical layer architecture and is classified into three different types depending on the histopathological phenotype (*48, 49*). Type I is characterized by isolated abnormal radial and/or tangential cortical lamination. Type II is the most prevalent form and characterized by dysmorphic neurons (subtype IIa) or additionally “balloon cells” (subtype IIb), which represent non-functional neural progenitor cells with enlarged cell bodies. Type III is associated with other pathologies such as hippocampal sclerosis, neoplasia, or vascular malformations. Ectopic neurons within the white matter can be found in all FCD types.

Diagnostic assessment of FCD in thin 2D histological sections suffers from several limitations. Firstly, the inherent focal nature of FCD makes it often difficult to identify pathological foci in large surgical specimens obtained from complete epileptic zone excision (*50–52*). This requires exhaustive and time-consuming serial sectioning. Secondly, suboptimal brain tissue orientation during FFPE block embedding and sectioning may impede accurate evaluation of cortical layering, which is crucial for FCD diagnosis. Lastly, FCD can manifest multifocally (*45*), rendering single thin sections inadequate for portraying the full pathological spectrum. The latter consideration is pivotal for estimating disease severity and devising appropriate follow-up interventions. Therefore, we were curious to assess whether 3D histology could help improve FCD diagnostics in difficult cases and accurately assess the extent of pathology in the entire surgical specimen. The compatibility of aDISCO with FFPE specimens from diagnostic archives allowed us to retrieve adequate numbers of historical specimens and benchmark 3D histology against traditional analyses.

We selected 7 FCD cases confirmed by 2D histology from the diagnostic archive of the University Hospital of Zurich (Table S1). All patients had undergone surgery due to refractory epilepsy. Only in three cases could the pathology be assessed throughout all cortical layers using conventional histology, due to the limitations of 2D techniques. For control, we selected 7 autopsy samples of age-matched patients with normal histology in the respective cerebral lobes from the diagnostic archive of the University Hospital of Zurich.

We stained neurons with anti-NeuN antibody to assess neuronal density across all cortical layers for 3D cortical profiling. Qualitative inspection of 3D stacks revealed focal distortion of the typical cortical layer architecture (cf. Fig. 6A), with only the molecular layer (I), the external granular layer (II), and the beginning of the white matter (WM) distinguishable, but not the layers in between (Fig. 7A and B). Furthermore, we observed ectopic neurons in the white matter. In contrast to the reference profile of normal cortex (cf. Fig. 6I), the cortical neuronal density profile in FCD exhibited only two peaks for layers I and II, while the other layers merged into a single hill that flattened slowly towards the white matter (Fig. 7C). This quantitative analysis demonstrates that 3D cortical profiling of neuronal density reflects the distortion of cortical layer architecture in FCD.

**Fig.7.**
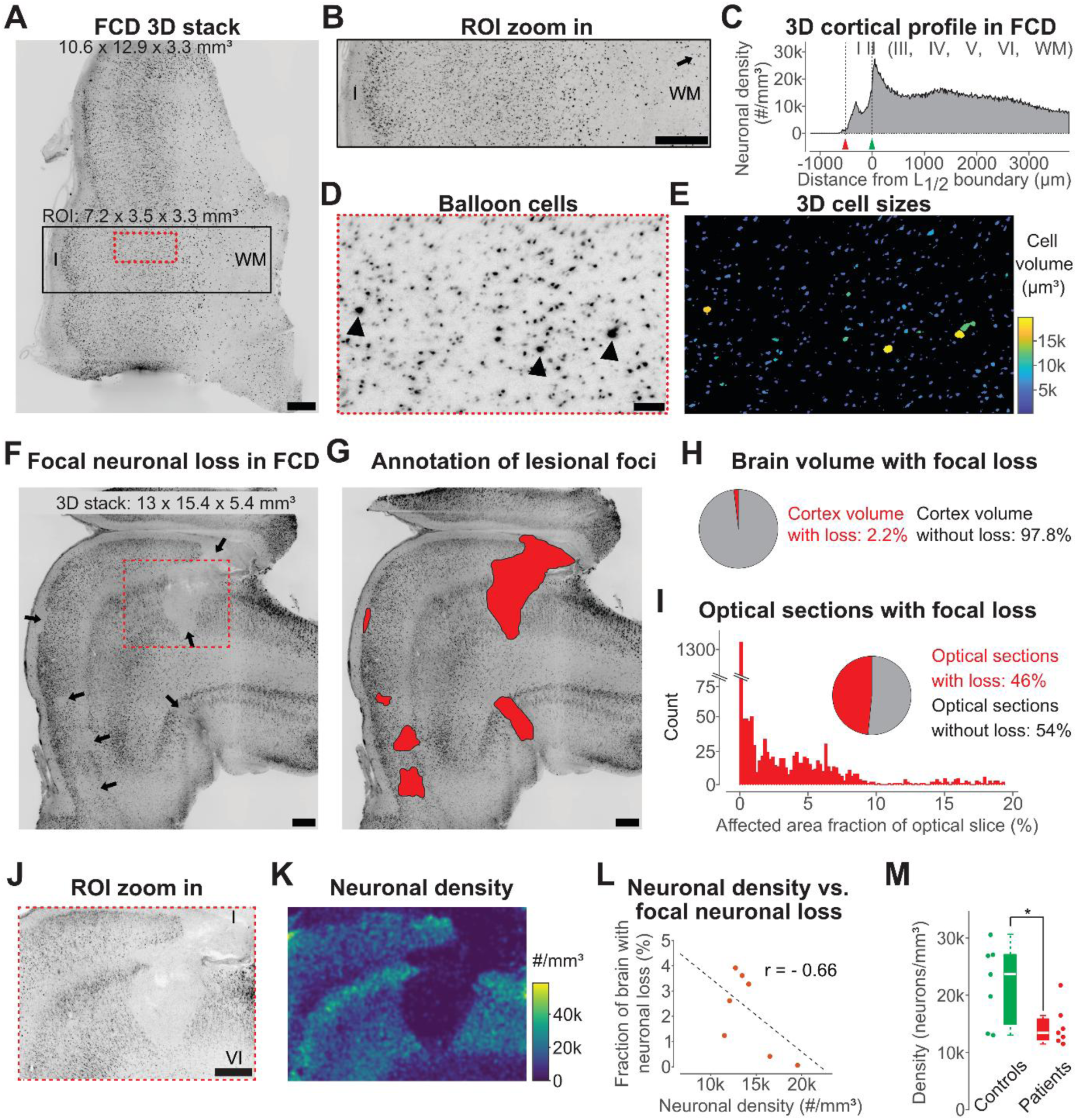
3D histological examination of FCD. (**A**) Stitched and 4x-downsampled 3D stack of archival human FCD cortex, imaged at 4x magnification using mesoSPIM V5. Distortion of layer architecture from the molecular layer (I) to the white matter (WM) is visible. Scale bar: 1 mm. (**B**) Region-of-interest (ROI) digital zoom into the black-lined box, highlighting distortion of cortical layer architecture and ectopic neurons in the WM (arrow). Scale bar: 1 mm. (**C**) Neuronal density profiling in FCD, displayed as number (#) of neurons per mm^3^ across all cortical layers, defined by the distance to the L1/2 boundary. Only two peaks, corresponding to layers I and II, can be distinguished from an otherwise undifferentiated hill which flattens into the WM. (**D**) ROI digital zoom into the red-dashed box, highlighting large, rounded balloon cells (arrowheads). Scale bar: 200 µm. (**E**) Automated detection of balloon cells based on their large cell size (yellow). Color scale indicates range of cell volume in µm^3^. (**F**) Stitched and 4x-downsampled 3D stack of archival human FCD cortex, imaged at 4x magnification using mesoSPIM V5. Multiple foci of neuronal loss (arrows) can be identified. Scale bar: 1 mm. (**G**) Illustration of 3D annotation of lesional foci. Scale bar: 1 mm. (**H**) Quantification of lesional foci in entire brain tissue blocks. (**I**) Quantification of lesional foci in optical sections. The histogram displays the count of optical sections in relation to the affected area per optical slice.(**J**) ROI digital zoom into the red-dashed box, highlighting a large focus of neuronal loss which affects all cortical layers I-VI. Scale bar: 500 µm. (**K**) Quantification of overall neuronal density in the lesional focus and the surrounding inconspicuous brain tissue, displayed as number (#) of neurons per mm^3^ in a heatmap. Color scale indicates cell number range in thousands (k). (**L**) Correlation of focal neuronal loss in percentage (%) and overall neuronal density displayed as number (#) of neurons in thousands (k) per mm^3^. Correlation coefficient r = −0.66. (**M**) Comparison of overall neuronal density quantification between FCD patients (red; n = 7) and controls (green; n = 7), displayed as number of neurons in thousands (k) per mm^3^ in box plots. Statistical analysis was performed by two-sided Student’s t-test. *p = 0.026.

To investigate an additional hallmark of FCD, we quantified the cell volume of cells. This analysis correctly detected the characteristic large, rounded balloon cells of FCD type IIb (Fig. 7D and E). Furthermore, during visual inspection of 3D stacks from FCD patients, we noticed multiple foci of neuronal loss, substantially varying in size and shape within and between patients (Fig. 7F). Based on the clinical history of FCD patients, these lesions were not caused by electrode implantations for two reasons: First, some patients who exhibited lesional foci had not undergone any electrode implantations. Second, for those who had been implanted with electrodes, the lesional foci did not match the electrode pattern in size and shape.

To analyze the extent of focal neuronal loss, we annotated all lesions in the entire 3D stacks (Fig. 7G). Quantitative volumetric analysis of neuronal loss foci demonstrated that these lesions affected only 2.2% of the cortical volume across FCD samples (Fig. 7H, Table S2). When we analyzed the 3D stacks as optical xy-sections along the z-axis, foci of neuronal loss were detected in less than half of optical sections (Fig. 7I). Most lesional foci were small and the maximally affected area per optical slice was 20%. Hence, the chance of missing such foci in thin sections, as in conventional 2D diagnostics, is high.

Finally, we quantified the overall neuronal density in brain volumes, including the lesional foci and the surrounding inconspicuous brain tissue in FCD (Fig. 7J and K). Bivariate analysis showed a strong negative correlation between overall neuronal density and the extent of lesional foci in FCD patients, indicating that FCD specimens with a higher fraction of brain volume with focal loss also exhibited overall lower neuronal density (Fig. 7L). Comparison between FCD and control groups revealed a significant reduction of neuronal density in FCD patients (median reduction of 43%; Fig. 7M, Table S2). Thus, the overall neuronal loss in FCD (43%; Fig. 7M) exceeded the loss caused by the lesional foci alone (2.2% of cortical volume; Fig. 7H), and affected brain tissue that would have been classified as healthy by conventional 2D histology. Therefore, quantitative 3D histology revealed additional neuronal loss undetectable by visual inspection.

In summary, our analyses demonstrate that aDISCO can be used to address specific pathologies by comprehensive and quantitative 3D histology, facilitated by a largely automated 3D data analysis pipeline.

## Discussion

Here, we introduce aDISCO, a method for 3D histology of whole-mount FFPE human tissue blocks, as a complementary diagnostic technique to conventional microscopy. We demonstrate that aDISCO achieves effective clearing and antibody staining throughout entire archival tissue blocks from different human organs and is compatible with a variety of antibodies, enabling specific immunolabeling of target structures down to the cellular level.

A central feature of aDISCO is the complete clearing and antibody staining of whole-mount FFPE human tissue blocks, whose sizes surpass the state-of-the-art for human FFPE specimens (*14, 19, 26, 28*) (Table 1). Imaging such large samples at cellular resolution with high throughput has only recently become a widely used approach due to the development of light-sheet microscopes with large working-distances (*29, 30, 53*). The largest human samples cleared and stained to date were freshly processed with the SHANEL protocol (*17, 18*). In this organic solvent-based tissue clearing method, the blood vessels of the respective human organs were perfused with chemical reagents to circumvent the diffusion barrier. With this procedure, a maximum imaging depth of 1 cm was reached for antibody-stained tissue. However, this approach is not applicable to archival human FFPE specimens from diagnostic archives, which contain large numbers of human samples from the distant past and thus constitute an untapped potential to study specific and rare diseases. The greatest imaging depth in FFPE tissue so far had been achieved with the organic solvent-based DIPCO protocol (*14*). However, DIPCO was applied to short-term formalin-fixed paraffin-embedded human biopsies (usually fixed < 24 hours) as opposed to long-term formalin-fixed paraffin-embedded human tissue specimens from autopsy (usually fixed > 24 hours up to three weeks, depending on organ type and size). In the DIPCO study, a maximum imaging depth of 1.6 mm was reached in human tumor biopsy specimens from six different human organs, characterized with five antibodies. To date, the only published human tissue clearing method performed on long-term formalin-fixed and paraffin-embedded human autopsy tissue was OPTIClear (*19*). With a combination of organic solvents used for deparaffination and a CLARITY-related clearing approach, this method reached a maximum imaging depth of 546 µm for antibody staining in human brain tissue. In contrast, in our study we reached a maximum imaging depth of 6.5 mm throughout the entire tissue block, with the sizes of the available samples being the limiting factor for the maximum imaging depth. The archival human FFPE tissue samples in our study had a maximum image size of 18.5 × 11.9 × 6.5 mm³ (lung sample) after clearing-induced tissue shrinkage of about 30% (*38*), i.e. with an original size on the cm-scale. The average sample size after clearing was 9.8 ± 4.2 mm × 7.6 ± 2.6 mm × 4.1 ± 1.0 mm³ (mean ± s.d. across 16 samples). Thus, with aDISCO we have established a tissue clearing method which enables access to whole-mount archival human tissue blocks by being compatible with both large human tissue specimens and samples from long-term FFPE storage (up to 15 years old in our study).

A second unique feature of aDISCO is its tested compatibility with 11 different human organs. This broad range of applications surpasses all existing clearing methods to date. Previous human tissue clearing methods were typically performed on one or a few tissue types (Table 1). Most studies focused on human brain cortex (*13, 15, 17, 19, 20, 22–27*), which is relatively straightforward to render transparent. In contrast, reports on dense tissues such as liver or skin, which are more challenging to clear (*9, 31*), are sparse. In human skin, previous work achieved a maximum imaging depth of 300 µm in immunolabeled tissue, while a depth of 1.4 mm was reached in unstained, autofluorescent tissue (*54, 55*). Similarly, in human liver most studies achieved imaging depths of only a few hundred micrometers in antibody-stained tissue (*56–59*). Notably, only two studies successfully imaged human liver volumes of 2.5 mm and 3 mm thickness, respectively (*60, 61*). However, human tissue clearing of archival FFPE skin or liver tissue had not been previously performed. With aDISCO, we achieved imaging depths of 3.4 mm in immunolabeled archival human skin and 3.5 mm in immunolabeled archival human liver tissue, values that again were only limited by the thickness of the available whole-mount FFPE blocks.

The third key advantage of aDISCO is its compatibility with at least 22 commonly used antibodies for cell typing, surpassing previous human tissue clearing studies. Antibody compatibility has been a significant challenge in tissue clearing and prior testing of antibodies with the specific chemicals and conditions used in clearing protocols has therefore been recommended (*10, 62*). Most antibodies were so far tested for compatibility only in non-archival human tissue (*17, 18, 25, 27*) (Table 1). The study with the largest (3 mm thick) FFPE human tissue specimens to date used staining with low-molecular-weight dyes (*16*), which, compared to antibodies, diffuse more easily into thick tissues, but lack the ability to differentiate distinct cell types. aDISCO overcomes these limitations, providing compatibility with diverse antibodies and complete immunolabeling throughout whole-mount archival human tissue blocks.

Finally, the ability to reparaffinize the cleared tissue blocks highlights the protocol’s versatility for subsequent conventional 2D histology, which is essential for backward compatibility and validation with the state-of-the-art method.

In a use-case study, we show that aDISCO enables standardized quantitative 3D histology of human samples collected from the diagnostic archive, and overcomes key limitations of conventional 2D diagnostics. In previous tissue clearing studies aiming to examine large human cortex specimens in 3D, staining was performed not with antibodies but with dyes that label all cells irrespective of distinct cell types (*16, 17*). Therefore, specific 3D quantification of neuron counts in large human cortex samples had not been performed to date. In our study, we demonstrate how aDISCO enabled the quantification of neuronal density and cortical layer profiling via specific immunolabeling of neurons and a largely automated data analysis pipeline. We applied this pipeline to a case series of focal cortical dysplasia (FCD), a neurodevelopmental malformation disorder characterized by subtle and focal pathology that often challenges diagnostic assessment by conventional 2D histology. 3D cortical profiling reflected the distortion of cortical layer architecture in FCD. Furthermore, cell volume analysis detected the large, rounded balloon cells characteristic of the most prevalent FCD subtype IIb, facilitating FCD diagnosis. Moreover, we observed multiple foci of neuronal loss, likely resulting from the long-lasting epilepsy with excitotoxic neuronal injury in neuronal assemblies. Previously, only two studies have reported neuronal loss in FCD using conventional 2D histology (*63, 64*). However, the extent of neuronal loss in the entire affected cortex had not been determined. aDISCO-based quantitative 3D histology revealed that the lesional foci constitute only a minor fraction of the cortical volume in whole-mount FFPE tissue blocks, a level of sparsity that conventional 2D histology is prone to overlook. Furthermore, there was additional neuronal loss not visible by eye and only detectable via quantification of neuronal cell densities. Together, these findings underscore the importance of quantitative 3D histology for holistic examination of human tissue specimens and promise to entail direct implications for diagnostic quality in FCD.

The primary current limitation of aDISCO is the relatively long, multistep protocol that prolongs processing time compared to conventional 2D histology. For this reason, aDISCO is best suited for retrospective 3D human histological studies and as a complementary diagnostic tool in particularly challenging cases, rather than for rapid routine histological diagnostics. Another consideration is the substantial data volume generated by high-magnification 3D light-sheet imaging of large human tissue blocks on the terabyte-scale. Managing, storing, and analyzing such data requires appropriate computational infrastructure and expertise. Lastly, a general limitation of human studies compared to genetically and environmentally controlled animal models is the considerable interindividual variability between patients. While this variability can complicate interpretation, statistically significant findings in human samples may hold greater translational relevance as they reflect authentic clinical heterogeneity.

In conclusion, human tissue clearing with aDISCO enables 3D histology across a broad range of organs and cell types in whole-mount archival human tissue blocks, previously inaccessible due to long-term FFPE storage. In combination with 3D data analysis, aDISCO allows quantitative 3D histology to address biological and pathological questions, including rare diseases, owing to the availability of human samples collected over years to decades. aDISCO’s broad applicability, compatibility with diverse antibodies, and integration with automated 3D data analysis can contribute to improved histological diagnostics and deeper insights into human diseases.

## Author contributions

A.M.R. designed the project, developed the protocol, conducted experiments, performed SPIM imaging, analyzed data, created movies, and wrote the manuscript with input from all co-authors. D.G. helped with data analysis, created movies, exchanged valuable ideas, and contributed to the manuscript write-up. M.C. provided shared technical help in conducting experiments and SPIM imaging. L.N. provided shared technical help in conducting experiments. L.F. established the stitching code and exchanged valuable ideas. F.F.V. and N.V. built the mesoSPIMs, designed the CAD figures, and provided continuous support together with P.B. R.R. wrote the ethical permit for this project with input by A.M.R. and exchanged valuable ideas. F.H. provided funding and expert input to the project. P.R. established the 3D data analysis pipeline, analyzed data, created movies, supervised the project, and contributed extensively to the figure design and manuscript write-up. A.A. provided the original idea, supervised the project, applied for funding, and contributed extensively to the manuscript write-up.

## Acknowledgements

We thank Irina Abakumova and Mirella Woodert for technical help with conventional 2D histology. We express our gratitude to José Maria Mateos Melero from the Center for Microscopy and Image Analysis, Zurich (ZMB) for technical assistance in light-sheet microscopy, and to André Wethmar for support in IT infrastructure. Furthermore, Florence Gillieron, Karen Sadoyama, and Magdalena Kloyer are kindly acknowledged for assistance in 3D image annotations. We thank Susanne Dettwiler for help in FFPE human sample selection from the diagnostic archive. Finally, we highly appreciate Viktor Koelzer’s expert input for figure design.

## Data and materials availability

All data needed to evaluate the conclusions in the paper are present in the paper and/or the Supplementary Materials.

## Materials and Methods

### Archival human FFPE samples

All procedures and the associated guidelines for processing archival FFPE human tissues were approved by the cantonal ethics committee Zurich (BASEC-No. 2022-00293). FFPE human tissue blocks were selected from the surgical and autopsy diagnostic archive at the Institute of Neuropathology and the Institute of Pathology and Molecular Pathology, University Hospital Zurich. The samples for testing compatibility of the aDISCO protocol with CNS regions and human organs originated from autopsies and were selected in a pseudonymized way. For the use-case study, FFPE brain surgical specimens from 7 FCD patients were chosen from the diagnostic archive. Patients with FCD were selected based on available clinical data and neuropathological diagnosis from conventional 2D histology. The hospital software database “KISIM” was used to check the medical history of FCD patients. Autopsy samples from 7 age-matched patients with normal histology in the respective cerebral lobes served as controls. For controls, only basic demographic information and brain location were available due to pseudonymization of control FFPE tissue blocks. Exclusion criteria for controls encompassed any pathologies in the respective brain region.

### Deparaffination of whole-mount archival human tissue blocks

FFPE human tissue blocks were melted for 1 hour at 60°C in a tissue embedding system TES99 (Medite). Samples were transferred into glass vials with closed top cap (C226-0020, Thermo Fisher Scientific). The remaining wax was removed by incubation in xylene (253-VL51TE, Thommen-Furler AG) 2 times each for 1 hour at 37°C, shaking at 50 rpm. Samples were rehydrated in decreasing serial solutions (100%, 95%, 90%, 80%, 70%, 50%, 25%) of ethanol (20821.321, VWR Chemicals) in Milli-Q® water (Merck Millipore) each for 1 hour at room temperature, shaking at 50 rpm, and finally in 1x PBS overnight at room temperature, shaking at 50 rpm.

### Dehydration and pre-clearing

Samples were dehydrated in increasing serial solutions (20%, 40%, 60%, 80%) of THF (186562, Sigma Aldrich) in Milli-Q® water each for 1 hour at room temperature, shaking at 50 rpm, followed by 2 times 100% THF each for 1 hour at room temperature, shaking at 50 rpm. Pre-clearing was performed in a 2:1 solution of DCM (270997, Sigma Aldrich) to THF overnight at room temperature, shaking at 50 rpm.

### Rehydration and bleaching

DCM was washed out by incubating the samples 2 times in 100% THF each for 30 minutes at room temperature, shaking at 50 rpm. Rehydration was achieved in decreasing serial solutions (80%, 60%, 40%) of THF in Milli-Q® water each for 1 hour at room temperature, shaking at 50 rpm. To pre-cool the samples for bleaching, the rehydration step in 20% THF in Milli-Q® water was performed for 1 hour at 4°C, shaking at 50 rpm. Bleaching was conducted in 28% H_2_O_2_ (1.08600.2500, Merck Millipore) with 4% THF in Milli-Q® water for 20 hours at 4°C in the dark, shaking at 50 rpm.

### Autofluorescence removal

To wash out the bleaching solution, samples were incubated 2 times in 1x PBS each for 1 hour at room temperature, shaking at 50 rpm. To remove remaining autofluorescence from highly pigmented samples, tissue blocks were incubated in 10 mM cupric sulfate (61230, Fluka) in 50 mM ammonium acetate (A-7330, Sigma Aldrich) buffer in Milli-Q® water, pH = 5, for 3 hours at room temperature in the dark, shaking at 50 rpm. The solution was washed out by subsequent incubation in 1x PBS 2 times each for 1 hour, and then overnight at room temperature, shaking at 50 rpm.

### Antigen retrieval

Depending on the respective antibodies, antigen retrieval was achieved by treating the samples with 10-30% formic acid (4724.3, Roth) solution in Milli-Q® water, pH = 2, for 3 hours at room temperature, shaking at 50 rpm. The solution was washed out by incubation in 1x PBS for 1 hour at room temperature, shaking at 50 rpm.

### Permeabilization and decolorization

Samples were first permeabilized with 0.2% TritonX-100 (X100, Sigma Aldrich) in 1x PBS 2 times each for 1 hour at room temperature, shaking at 50 rpm. Stronger permeabilization and decolorization was achieved by subsequent incubation in 25% urea (U4884, Sigma Aldrich) + 25% NNNN-tetrakis(2-OH-propyl)ethylenediamine (122262, Sigma Aldrich) + 15% TritonX-100 in 1x PBS overnight at 37°C, shaking at 50 rpm. Permeabilization was continued in 0.2 % TritonX-100 + 2.3% glycine (G8898, Sigma Aldrich) + 20% DMSO (D8418, Sigma Aldrich) + 0.5 mM methyl-beta-cyclodextrin (332615, Sigma Aldrich) + 0.2% trans-1-acetyl-4-OH-L-proline (441562, Sigma Aldrich) + 0.1% sodium azide (S8032, Sigma Aldrich) in 1x PBS for 3 days at 37°C, shaking at 50 rpm.

### Blocking and immunolabeling

For blocking and antibody staining, samples were transferred from glass vials into 6-(92006, TPP) or 12-well plates (92012, TPP), depending on sample size, and sealed with parafilm (10018130, Bemis) during incubation to avoid evaporation. Blocking was performed in 0.2% Tween-20 (P2287, Sigma Aldrich) + 0.1% heparin (10 mg/ml, 2835358, B. Braun Medical AG) + 5% DMSO + 6% donkey serum (017-000-121, Jackson Immuno Research) in 1x PBS for 2 days at 37°C, shaking at 50 rpm. Immunolabeling was conducted in 0.2% Tween-20 + 0.1% heparin (10 mg/ml) + 5% DMSO + 0.1% sodium azide (staining buffer) for up to 4 weeks, depending on sample size and antibody diffusion capacity, at 37°C, shaking at 50 rpm. To enhance the diffusion capacity of antibodies, immunolabeling was carried out gradually, starting with low antibody concentrations and adding more and more antibodies to the staining solution to finally reach the calculated final antibody concentrations at the end of the staining time. For this gradual immunolabeling step, the starting and final antibody concentrations as well as the staining times were adapted to the specific antibodies used in this study (Table S3). For example, when the final dilution of an antibody with a stock concentration of 1.5 mg/ml was 1:200 and the staining time was two weeks, we started with a dilution of 1:800 and added the same dilution another three times every half a week, thus reaching the final dilution of 1:200 after two weeks.

Between primary and secondary antibody staining, samples were washed in staining buffer 2 times each for 1 hour, then 2 times each for 2 hours, and then overnight at room temperature, shaking at 50 rpm. After secondary antibody incubation, samples were washed in staining buffer 2 times each for 1 hour, then 2 times each for 2 hours, and then for 2 days at room temperature, shaking at 50 rpm.

### Dehydration and clearing

Samples were dehydrated in increasing serial solutions (20%, 40%, 60%, 80%) of THF in Milli-Q® water each for 1 hour at room temperature, shaking at 50 rpm, followed by 2 times 100% THF each for 1 hour at room temperature, shaking at 50 rpm. Final delipidation was performed in a 2:1 solution of DCM to THF overnight at room temperature, shaking at 50 rpm, followed by incubation in 100% DCM 2 times each for 2 hours at room temperature, shaking at 50 rpm. Refractive index (*n*_D_ = 1.56) matching was achieved in DBE (108014, Sigma Aldrich) overnight at room temperature, non-shaking.

### Reparaffination of cleared tissue blocks

DBE was washed out by incubating the tissue blocks in 100% methanol (203-VL03K-/1, Thommen-Furler AG) 2 times each for 1 hour at room temperature, shaking at 50 rpm. Samples were rehydrated in decreasing serial solutions (80%, 60%, 40%, 20%) of methanol in Milli-Q® water each for 1 hour at room temperature, shaking at 50 rpm, followed by incubation in 1x PBS overnight at room temperature, shaking at 50 rpm. Tissue blocks were again dehydrated in increasing serial solutions (25%, 50%, 70%) of ethanol in Milli-Q® water each for 1 hour at room temperature, shaking at 50 rpm. Then, samples were put in the Leica ASP300S Fully Enclosed Tissue Processor (Leica Biosystems) and the following program started: 100% ethanol for 45 minutes, 1 hour, and 30 minutes at 37°C, followed by 100% ethanol 2 times each for 1 hour at 45°C; then xylene for 45 minutes and 2 times each for 1 hour at 37°C; and finally paraffin (39601006, Leica Biosystems) for 1 hour and 15 minutes, 1 hour and 30 minutes, and 2 hours at 62°C.

### 2D histology

2-µm thick tissue sections were cut from the tissue blocks, using a semi-automated rotary microtome, (RM2245, Leica Biosystems) and mounted onto slides (0810531, HistoBond®). Slides were deparaffinized by incubation in xylene 2 times each for 10 minutes at room temperature. Slides were rehydrated in decreasing serial solutions (100%, 95%, 90%, 80%, 70%) of ethanol in Milli-Q® water each for 10 minutes at room temperature, followed by pure Milli-Q® water for 10 minutes at room temperature.

For H&E staining, sections were first incubated in filtered hematoxylin Harris (2E-056, Waldeck) for 5 minutes at room temperature, followed by rinsing in Milli-Q® water. Sections were differentiated in 0.185% HCl (124630010, Acros Organics) in Milli-Q® water by 4 quick immersions at room temperature, followed by rinsing in tap water. Bluing was performed in 0.015% ammonia water (1.05432.1000, Merck Millipore) by 6 quick immersions at room temperature, followed by rinsing in Milli-Q® water and 4 quick immersions in 70% ethanol at room temperature. Sections were incubated in 0.075% eosin (1B-425, Waldeck) + 0.0075% phloxin (15926.0025, Merck Millipore) + 0.5% acetic acid (1.00063.1011, Merck Millipore) + 84% ethanol in Milli-Q® water for 3 minutes at room temperature. After rinsing in 100% ethanol and xylene, they were finally covered with coverslips (01012227, Biosystems).

For antibody staining, a Ventana Bench Mark Ultra Plus fully automated staining machine (Roche) was used, containing the Bench Mark Ultra LCS staining kit. Staining was visualized with HRP DAB included in the kit. Slides were scanned using a Hamamatsu C9600 digital slide scanner (Hamamatsu Photonics).

### 3D light-sheet imaging

Light-sheet imaging was performed using a custom-built mesoSPIM V5 (*29*). For all image settings, the following lasers and filter combinations were applied: 488 nm laser with a 520/35 BrightLine HC filter (AHF analysentechnik AG, part number #F37-520SG) or QuadLine Rejectionband ZET405/488/561/640 filter (AHF analysentechnik AG, part number #F57-405SG); 561 nm laser with 561 Razor Edge LP filter (AHF analysentechnik AG, part number #F76-561) or QuadLine Rejectionband ZET405/488/561/640 filter; 640 nm laser with 647 Razor Edge LP Filter (AHF analysentechnik AG, part number #F76-647); 647 nm laser with QuadLine Rejectionband ZET405/488/561/640 filter. Laser intensities were individually set for the samples/each experiment to achieve the best SNR. For instance, for the autofluorescence removal experiment the same settings were used for all samples. For overview whole-mount 3D images, the 1x objective was used at zoom levels of 0.63x, 1x or 2x, depending on sample size. Samples were imaged in a 40×40×40 mm cuvette filled with DBE (*n_D_* = 1.56) and z-step size was chosen to reach (near) isotropic voxel size. Camera exposure time was 10 ms.

For the use-case study, tile scanning of each entire cortex sample was performed with the 1x objective at 4x zoom. Samples were imaged in a 30×30×30 mm cuvette filled with DBE with a z-step size of 1 µm. Camera exposure time was 10 ms.

For high-magnification ROI imaging of almost all samples, a 2x objective (MVPLAPO2XC, Olympus/Evident) was used in combination with a dipping cap (205915, LaVision BioTec). The front cover glass of the dipping cap was removed and replaced with a 40×40×40 mm cuvette (Portmann Instruments) filled with DBE. This objective was used at zoom levels of 2x, 4x or 6.3x, and z-step size was chosen to reach (near) isotropic voxel size. With this system at highest zoom level (6.3x) and an NA of ∼0.2, the standard Rayleigh criterion for the lateral (xy) diffraction-limited resolution at 561 nm wavelength is:

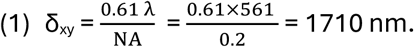

Camera exposure time was 10 ms.

Alternatively, for the peripheral nerve sample a modified detection path based on the Benchtop mesoSPIM (mesoSPIM V6) with a Mitutoyo objective turret (Mitutoyo) and a Mitutoyo G Plan Apo 20x objective was used (*30*). Samples were imaged in a 40×40×40 mm cuvette filled with DBE (*n_D_* = 1.56) and z-step size was chosen to reach (near) isotropic voxel size. Camera exposure time was 20 ms.

### Qualitative image processing

For 3D data analysis, an AMD Ryzen W137 Threadripper PRO Workstation with 2x NVIDIA Geforce RTX 3090 24GB GDDR6X graphic cards (Brentford) was used. Fiji software (64 bit version 2.3.0/1.53f51 (*65*)) was employed for qualitative image processing, containing cropping, contrast adjustment, background subtraction, contrast limited adaptive histogram equalization (CLAHE), color and look up table (LUT) intensity assignment, channel merging, generation of hyperstacks with temporal color code, generation of substacks and maximum intensity projections, generation of movies, and scale bar assignments. The volume viewer plugin (*66*) was applied for volume rendering. The 3DScript plugin (*67*) was used for volume rendering and for generating complex movie sequences via encoded movie scripts. Transparency was altered by adapting the alpha-value. Adobe Premiere Elements (Adobe version 2021) was applied for further image and movie processing, consisting of sharpening, brightening, movie clipping and text labeling.

### Quality assessment of light-sheet imaging data

#### Fourier ring correlation quality estimate (FRC-QE)

FRC-QE was conducted using a published Fiji package (*36*) with the default parameters.

#### Dendritic cross-sectional profiles

Dendrites with small diameter were visually selected and a cross-sectional profile of 5-10 pixel width drawn using the *straight line* tool in Fiji. Values were saved in a csv-file and imported to MATLAB (MathWorks, version 2022b) for Gaussian fitting with the *cftool* toolbox. The full width of half maximum (FWHM) was calculated from the standard deviation of the fitted Gaussian.

#### Tracing of neuronal processes

Main processes of example dopaminergic neurons were traced using the *brush* tool of the tracing framework webKnossos (*37*) in the web interface.

#### Mean fluorescence intensity (MFI) projection and z-profile

Codes for the assessment of MFI projection and z-profile were written in MATLAB. For both calculations, the background of the sample was excluded by setting all background pixels without sample to not-a-number (NaN). The MFI projection was calculated as the mean projection value along the z-axis and plotted as heatmap in xy-direction. The z-profile was calculated as the mean fluorescence intensity for each imaging plane (in z) across the sample, with the standard error of the mean (s.e.m.) computed across the pixels of each plane.

### Stitching

3D image tiles were automatically stitched via a custom-written code in R studio (version 1.2.1335, R version 3.6.0 (*68*)), using Fiji and TeraStitcher (version 1.11.6. (*69*)).

### Neuronal cell segmentation and detection

To automatically and reliably detect neurons in human brain cortex, a three-step procedure was implemented in Python (versions 2.7. and 3.10.12 (*70, 71*)). First, stitched raw stacks were segmented using a supervised random forest machine learning classifier (ILASTIK version 1.4.0rc6 (*43*)) into coherent 3D particles that were then identified as initial neuronal cell candidates using an established cell detection pipeline (ClearMap 1.0 (*44*)). To train the ILASTIK classifier, ground truth was manually annotated volumetrically as foreground vs. background in 13 representative substacks of 200 x 200 x 200 voxels. Representative substacks were generated from the entire 15 TB-sized data set, representing various regions (transitions of different cortical layers and white matter, regions with high and lower SNR) from different samples (controls and FCD patients). To fit the input into the memory for ILASTIK processing, the ClearMap pipeline was parallelized. Segmentation was performed such that overdetection was more likely than underdetection to enable subsequent refinement by discarding false positives. Refinement of cell candidates is conceptually similar to an existing pipeline (*72*) for murine cell detection, but has been implemented independently here for our human data. Initial neuronal cell candidates were detected based on local intensity maxima using ClearMap. For quality checks, results were visualized in Fiji. Second, initial cell candidates were merged using a custom-written 3D-peak finder based on 3D gradient ascent. Briefly, for each initial neuronal cell candidate determined by ClearMap in step 1, the local raw 3D image (31 x 31 x 91 voxels in xyz) was extracted and smoothed with a Gaussian filter (standard deviation of 1.0, 1.0, and 2.5 pixels in x-, y-, and z-direction, respectively). Within this local environment, the position of the detected cell was iteratively optimized to follow the maximum image intensity until the algorithm converged to a local maximum. Upon this gradient-based refinement of neuron positions, cells that ended up at the same maximum were merged. Code for gradient-based refinement and merging of neuronal cell candidates is available on Github: https://github.com/PTRRupprecht/Cell_Detection/tree/main/Gradient_cell_merger. Third, the refined neuronal cell candidates were classified as cells or non-cells using a DCNN-based binary classifier. This procedure served to exclude non-neuronal somata bright structures such as proximal axons, hot pixels of the camera sensor, or spurious noise. Briefly, each 100 neuronal cell candidates as output from the second processing step were chosen at random from 19 different 3D samples, resulting in a total of 1900 neuronal cell candidates for ground truth annotation and validation. These neuronal cell candidates were manually annotated by visually inspecting xy-sections (31 x 31 pixels) and xz-sections (31 x 91 pixels), centered at the location of the candidate cell. Each neuronal cell candidate was assigned either a “cell” or “non-cell” label. A Python-based interactive tool used to perform these annotations is available on Github: https://github.com/PTRRupprecht/Cell_Detection/tree/main/Label_tool. Next, DCNNs were trained based on the annotated ground truth to predict whether a detected neuronal cell candidate was a cell. Three networks for different sections (xy, xz, yz) were combined by adding their outputs. All networks were standard convolutional networks with two 2D convolutional layers with ReLu activation units, followed each by a 2 x 2 max pooling layer, a subsequent set of two dense layers with ReLu activation units, and a final layer with 2 neurons. These two output neurons represented the two classes (“cell” and “non-cell”). The network was defined in PyTorch (*73*) and trained with a standard Adagrad (*74*) optimizer, using cross entropy loss with a batch size of 10. The classification performance plateaued after approximately 1500 training epochs (Fig. S5C). 1800 randomly selected neuronal cell candidates were used for training. Performance was evaluated using cross-validation with randomly selected withheld neuronal cell candidates (100 random cell candidates withheld of a total of 1900). To improve cell classification, averaging across deep network ensembles was used (*75*). Five instantiations of each network type (xy, xz, yz) were trained and their predictions were linearly combined for ensemble averaging across 15 models. Based on ensemble predictions, neuronal cell candidates were accepted or discarded. In cross-validated performance evaluation, the classifier was correct for 92.2% of neuronal cell candidates and incorrect for 7.8%, with false positives and false negatives at 4.2% and 3.6%, respectively. Python code to train and apply the neuronal cell candidate classifier together with an example ground truth data set of manually annotated neuronal cell candidates is available on Github: https://github.com/PTRRupprecht/Cell_Detection/tree/main/Binary_classification_neuron_candidates.

### Quantification of neuronal density

To quantify neuronal density, neurons in the 3D stack were detected as described above, yielding the total number of detected neurons. Subsequently, the number of detected neurons was divided by the volume occupied by the brain sample. The volume of the brain sample was obtained by a binary 3D mask computed in Python, using a manually defined global threshold. The resulting 3D mask was corrected and refined by manual annotation of the 12x-downsampled segmented stack in 3D with three interactive views (xy, xz, yz) using ITK-SNAP (version 3.6.0 (*76*)). To distinguish cortical vs. white matter brain volumes, cortical regions of 12x-downsampled imaging stacks were manually annotated in 3D with three interactive views (xy, xz, yz) using ITK-SNAP (version 3.6.0). Manual annotation of focal neuronal loss regions was performed in ITK-SNAP (version 4.0.1) by carefully inspecting the 4x-downsampled brain tissue blocks in 3D with three interactive views (xy, xz, yz).

### Neuronal cell density profiles across cortical layers

To assign “cortical depth” normalized to each position in a stack by the local cortical thickness, landmarks within the cortical sample were annotated, hyperplane fits of these annotations applied, and the distance between these hyperplanes scaled to normalize for cortical thickness. First, to annotate landmarks, we identified the outer brain boundary, the boundary between the cortical layers 1 and 2, the stripe of increased neuronal density in layer 4, and the boundary between layer 6 and the white matter. Xy-sections were manually annotated with 5-8 coordinates for the respective landmark in a custom-written interactive Python script. Every 100 pixels (100 µm) in the z-direction, an xy-section was annotated and a 3^rd^ order polynomial hyperplane was fitted to each of the landmark layers. For each neuron in the 3D volume, the shortest distance to each of the landmark hyperplanes was determined, which enabled the computation of the local relative distance between two landmark layers.

### Figure design

Figures were designed in Affinity Designer 2 (version 2.3.1) and Adobe Illustrator (version 2025). CAD models were designed in Autodesk Inventor (version 2023).

## Supplementary Material

### Supplementary Figures

**Fig. S1.**
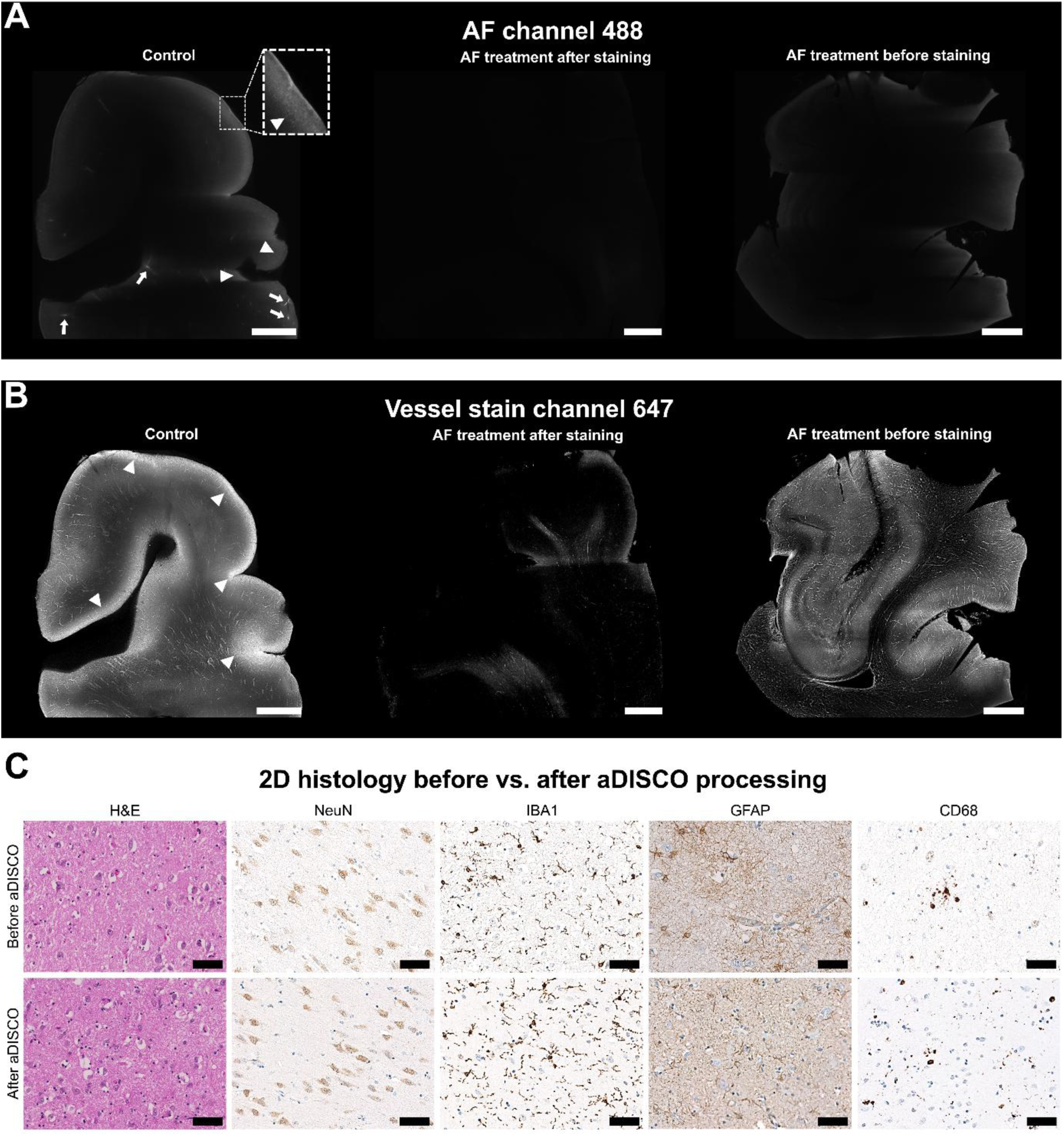
Autofluorescence (AF) removal and reparaffination. **(A)** Unstained AF channel (488 nm) of archival human brain samples shows autofluorescent artifacts in the control sample, arising from vessels with coagulated blood (arrows) and lipofuscin (arrowheads) despite bleaching with 28% hydrogen peroxide. AF treatment applied either before or after staining removes these artifacts. **(B)** Vessel channel (647 nm) shows CD13-immunolabeled vasculature. In the control sample, lipofuscin autofluorescence interferes with the vessel signal (arrowheads). AF treatment after staining removes these artifacts but also substantially reduces the vessel signal, whereas treatment before staining preserves it. All samples (A-B) were imaged at 0.63x magnification using mesoSPIM V5 and processed with identical settings. Scale bars: 2 mm. (**C**) Conventional 2D histology before vs. after aDISCO processing. 2-µm thin sections cut from the same human brain FFPE tissue block before and after performing the entire aDISCO protocol with reparaffination after tissue clearing. Sections were stained for hematoxylin and eosin (H&E) as well as for different antibodies, and imaged using a Hamamatsu C9600 digital slide scanner. Scale bars: 50 µm.

**Fig. S2.**
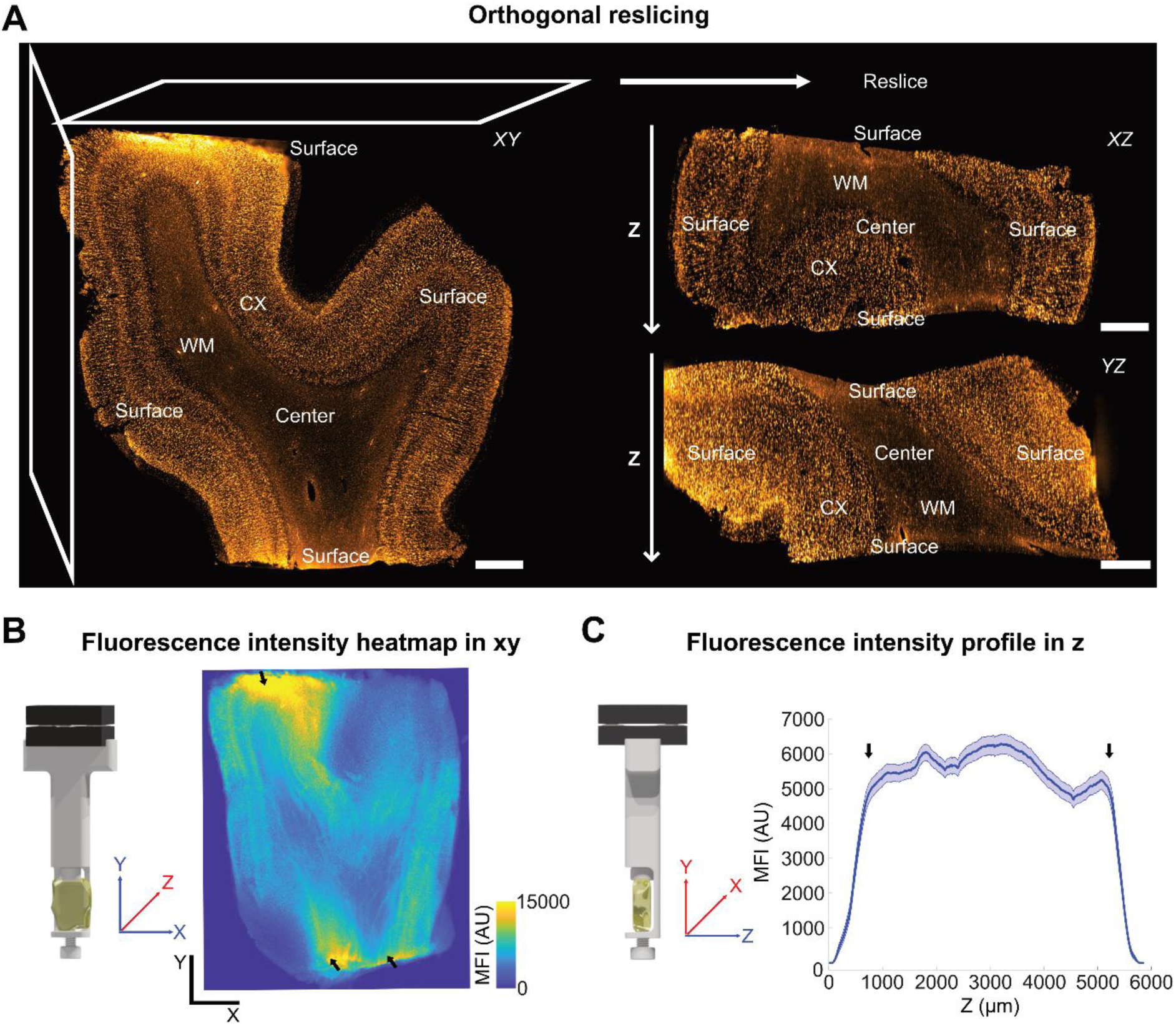
Effective clearing and immunolabeling of aDISCO-processed archival human brain cortex. (**A**) Large archival human brain cortex sample, immunolabeled for neurons via anti-NeuN antibody and imaged at 1.25x magnification using mesoSPIM V5. Left: Slice in xy-projection from the sample center in z-direction. Right: Slice in xz-(top) and yz-projection (bottom) from the sample center in y- and x-direction, respectively, generated via reslicing of the 3D stack. In all three projections, continuous staining signals in the cortex (CX) from the surfaces to the center as well as effective (non-blurry) clearing of the entire sample, including the neuron-poor and lipid-rich white matter (WM), can be observed. This continuity through the entire sample thickness is particularly visible in the resliced xz-(top) and yz-projection (bottom) along the z-axis (z arrows). Scale bars: 1 mm. (**B**) Mean fluorescence intensity (MFI) projection averaged over all planes in z (red axis) and plotted as heatmap in xy (blue axes), as illustrated by the computer-aided design (CAD) model. Focal high-intensity pixels at the top and bottom are due to screwing of the sample to the holder for imaging (black arrows). Color scale indicates MFI range in arbitrary units (AU). Scale bars: 2 mm. (**C**) Z-profile of mean fluorescence intensity (MFI) in arbitrary units (AU), averaged over all xy-pixels (red axes) per imaging plane and plotted along the z-axis (blue) with s.e.m. (light blue), as illustrated by the computer-aided design (CAD) model. Arrows indicate the beginning and end of the sample.

**Fig. S3.**
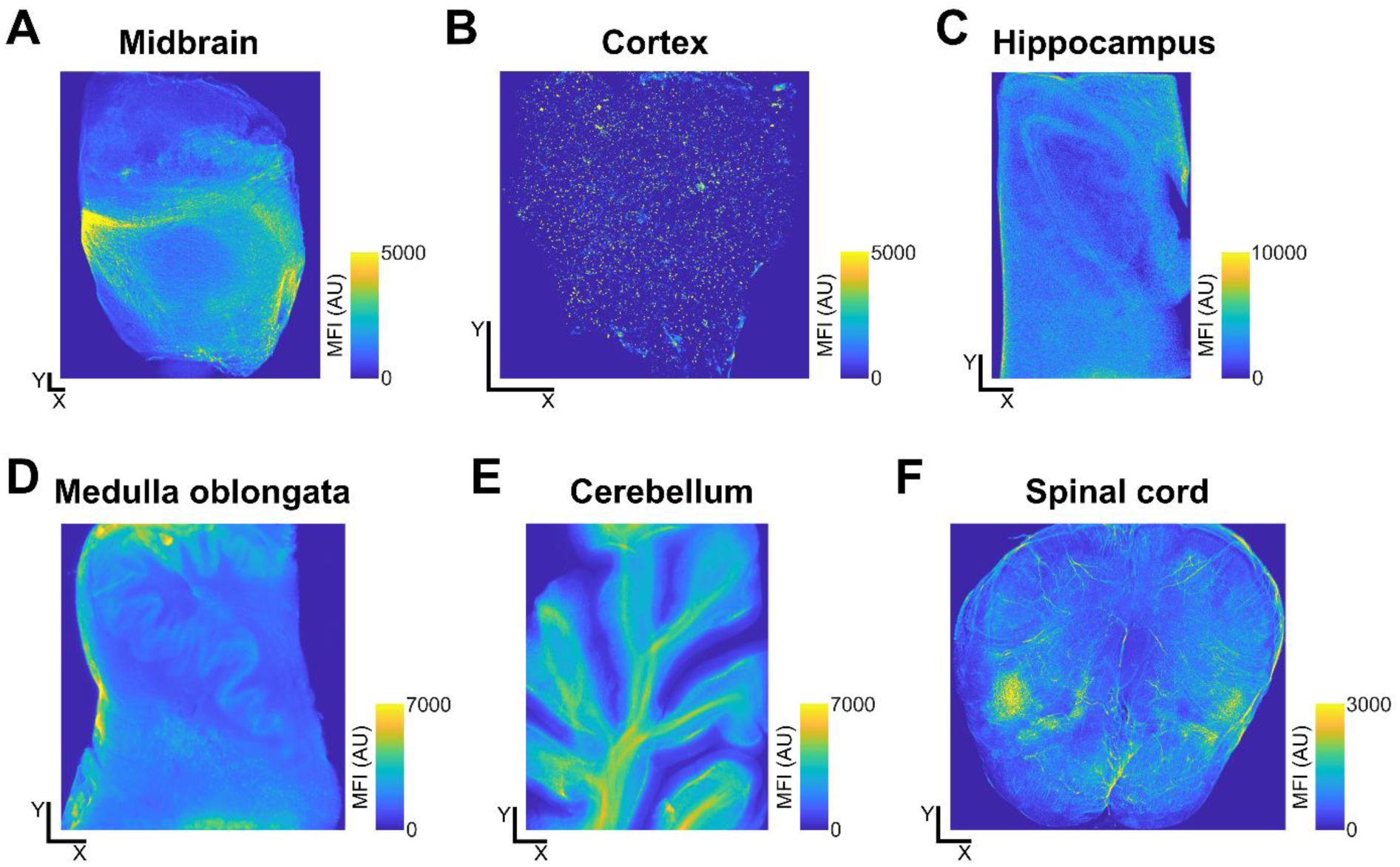
Mean fluorescence intensity (MFI) projections of different archival human CNS regions. (**A-F**) MFI projections averaged over all planes in z and plotted as heatmaps in xy. Color scales indicate MFI ranges in arbitrary units (AU). Scale bars: 1 mm.

**Fig. S4.**
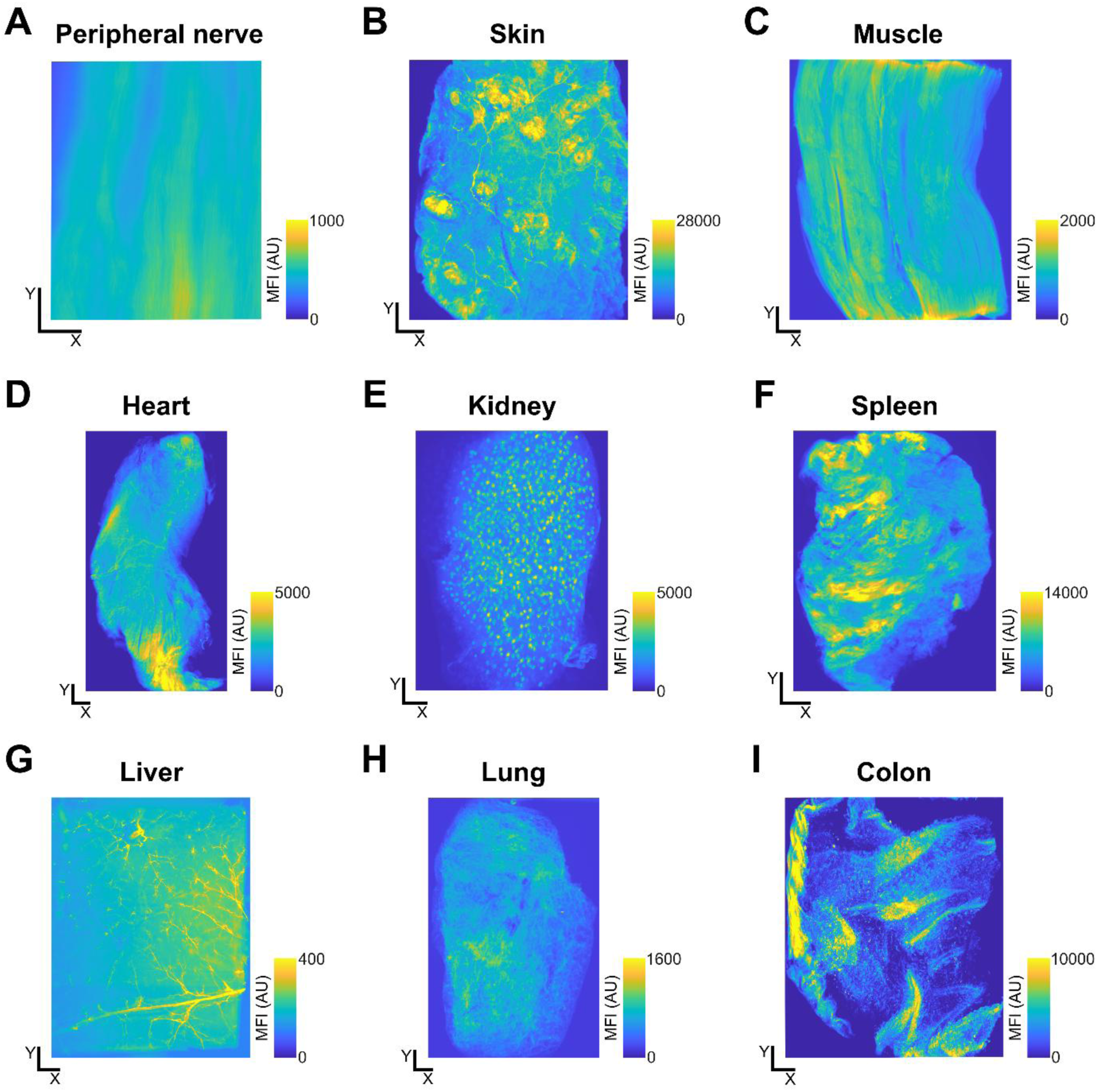
Mean fluorescence intensity (MFI) projections of different archival human organs. (**A-I**) MFI projections averaged over all planes in z and plotted as heatmaps in xy. Color scales indicate MFI ranges in arbitrary units (AU). Scale bars: 1 mm.

**Fig. S5.**
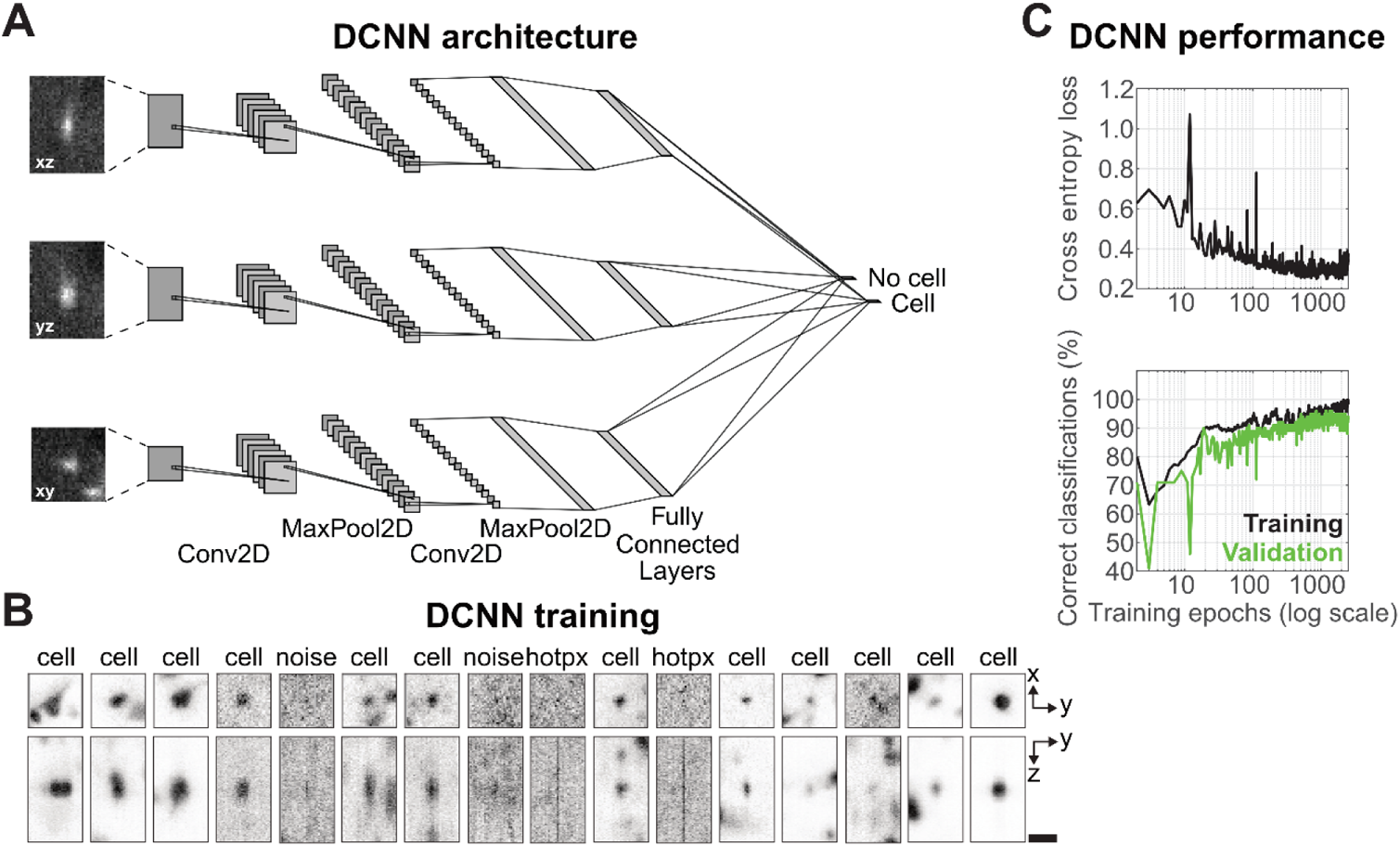
DCNN architecture and performance. (**A**) DCNN architecture for all three projections xz, yz, and xy. Each network consists of two convolutional (conv) 2D layers with ReLu functions, followed by a 2 x 2 max pooling (pool) 2D layer, two subsequent fully connected layers with ReLu functions, and a final layer with two neurons representing “cell” vs. “non-cell”. The output of the three networks is combined to decide whether a cell candidate is a “cell” or “non-cell”. (**B**) DCNN ground truth examples of the training input in different projections to classify “cell” vs. “non-cell” such as noise or hot pixels (hotpx) from the recording camera. For all excerpts, the full side lengths are 46.5 µm x 46.5 µm x 91 µm. Scale bar: 25 µm. (**C**) DCNN performance. Top: The cross entropy loss function decays during training and reaches a plateau after 1500 training epochs. Bottom: The percentage of correct classifications increases both for the training set (black) and for the validation set (green), and reaches a plateau of > 90% after 1500 training epochs.

### Supplementary Tables

**Table S1.**
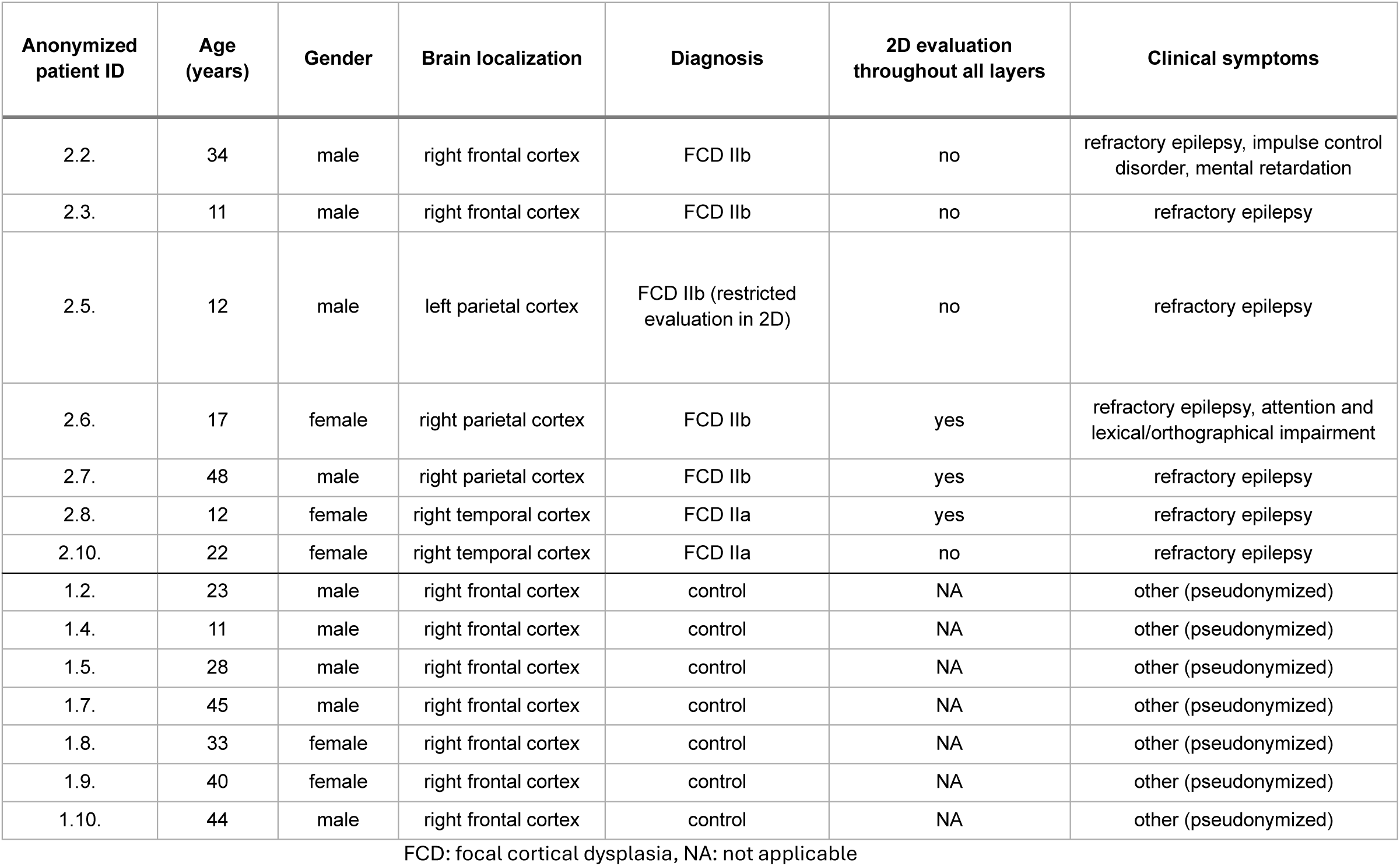
| Characteristics of FCD patients and controls.

| Anonymized patient ID | Age (years) | Gender | Brain localization | Diagnosis | 2D evaluation throughout all layers | Clinical symptoms |
| --- | --- | --- | --- | --- | --- | --- |
| 2.2. | 34 | male | right frontal cortex | FCD IIb | no | refractory epilepsy, impulse control disorder, mental retardation |
| 2.3. | 11 | male | right frontal cortex | FCD IIb | no | refractory epilepsy |
| 2.5. | 12 | male | left parietal cortex | FCD IIb (restricted evaluation in 2D) | no | refractory epilepsy |
| 2.6. | 17 | female | right parietal cortex | FCD IIb | yes | refractory epilepsy, attention and lexical/orthographical impairment |
| 2.7. | 48 | male | right parietal cortex | FCD IIb | yes | refractory epilepsy |
| 2.8. | 12 | female | right temporal cortex | FCD IIa | yes | refractory epilepsy |
| 2.10. | 22 | female | right temporal cortex | FCD IIa | no | refractory epilepsy |
| 1.2. | 23 | male | right frontal cortex | control | NA | other (pseudonymized) |
| 1.4. | 11 | male | right frontal cortex | control | NA | other (pseudonymized) |
| 1.5. | 28 | male | right frontal cortex | control | NA | other (pseudonymized) |
| 1.7. | 45 | male | right frontal cortex | control | NA | other (pseudonymized) |
| 1.8. | 33 | female | right frontal cortex | control | NA | other (pseudonymized) |
| 1.9. | 40 | female | right frontal cortex | control | NA | other (pseudonymized) |
| 1.10. | 44 | male | right frontal cortex | control | NA | other (pseudonymized) |

FCD: focal cortical dysplasia, NA: not applicable

**Table S2.** | Summary of neuronal loss analyses in distinct FCD patients.

| **Patient ID** | **#2** | **#3** | **#5** | **#6** | **#7** | **#8** | **#10** |
| --- | --- | --- | --- | --- | --- | --- | --- |
| Number of visible lesional foci | 1 | 2 | 11 | 25 | 2 | 5 | 16 |
| Affected volume of brain (%) | 1.25 | 0.41 | 2.65 | 3.67 | 0.06 | 3.98 | 3.32 |
| Affected volume of cortex (%) | 1.46 | 0.46 | 3.05 | 3.77 | 0.08 | 4.43 | 4.02 |
| Affected optical slices (%) | 4.7 | 48.1 | 89.2 | 54.4 | 14.8 | 53.9 | 75.2 |
| Mean affected area fraction per optical slice (%) | 0.73 | 0.27 | 4.00 | 2.47 | 0.05 | 2.73 | 5.02 |
| Cell density (1000 per mm<sup>3</sup>) | 11.5 | 16.5 | 12.1 | 13.5 | 21.7 | 12.7 | 14.2 |
| Cell density change compared to median density of control<br>(23.7 x 1000 per mm<sup>3</sup>) | -51.3% | -30.3% | -48.9% | -43.1% | -8.0% | -46.1% | -40.1% |

**Table S3.** | Antibodies list.

| Antibody (species, clonality) | Source (order number) | LOT number | RRID | Dilution (start → final) | Staining time (3D histology) |
| --- | --- | --- | --- | --- | --- |
| AlexaFluor® 647 anti-Myelin Basic Protein (mouse, monoclonal) | Biologend (808408) | B322585 | AB_2814552 | 1:200 → 100 | 5 days |
| Anti-Aminopeptidase N/CD13 (goat, polyclonal) | R&D systems (AF2335) | VRR0320081 | AB_2227288 | 1:400 → 100 | 2 weeks |
| Anti-α-Actin, smooth muscle-Cy3™ (mouse, monoclonal) | Sigma Aldrich (C6198) | 86029 | AB_476856 | 1:800 → 200 | 2 weeks |
| Anti-Collagen IV (goat, polyclonal) | Abcam (ab193951) | GR274637-5 | AB_94799 | 1:1000 → 250 | 2 weeks |
| Anti-E Cadherin (rabbit, monoclonal) | Cell Signaling (3195S) | 15 | AB_2291471 | 1:800 → 200 | 2 weeks |
| Anti-GFAP (rabbit, polyclonal) | Agilent (Z0334) | 00051304 | AB_10013382 | 2D: 1:13000 | - |
| Anti-Human CD45 (mouse, monoclonal) | Agilent (M0701) | 41405417 | AB_2314143 | 1:800 → 200 | 2 weeks |
| Anti-Human Nephrin (sheep, polyclonal) | R&D systems (AF4269) | ZMU0221031 | AB_2154851 | 1:800 → 200 | 2 weeks |
| Anti-Human Podocalyxin (goat, polyclonal) | R&D systems (AF1658) | JLD0221051 | AB_354920 | 1:400 → 100 | 2 weeks |
| Anti-IBA1 (rabbit, polyclonal) | Fujifilm Wako (019-19741) | LER0547 | AB_839504 | 1:800 → 200<br>2D: 1:1000 | 1 week |
| Anti-Laminin (rabbit, polyclonal) | Sigma Aldrich (L9393) | VG3038968 | AB_477163 | 1:800 → 200 | 2 weeks |
| Anti-N Cadherin (rabbit, polyclonal) | Abcam (ab18203) | GR3404600-2 | AB_444317 | 1:200 → 50 | 2 weeks |
| Anti-NeuN (rabbit, polyclonal) | Merck Millipore (ABN78) | 3692639 | AB_10807945 | 1:800 → 200<br>2D: 1:400 | 2 weeks |
| Anti-Neurofilament pan-neuronal (mouse, monoclonal) | Biologend (837801) | B279610 | AB_2565383 | 1:800 → 200 | 2 weeks |
| Anti-Neurofilament M (rabbit, polyclonal) | Biologend (841001) | B248163 | AB_2565457 | 1:800 → 200 | 5 days |
| Anti-Parvalbumin (rabbit, polyclonal) | Swant (PV27) | PV27a_221215_200 | AB_2631173 | 1:1000 → 250 | 2 weeks |
| Anti-PGP9.5 (rabbit, polyclonal) | Agilent (Z5116) | 20074146 | AB_2622233 | 1:400 → 100 | 2 weeks |
| Anti-Tryptophane Hydroxylase (mouse, monoclonal) | Merck Millipore (MAB5278) | 3786041 | AB_2207684 | 1:800 → 200 | 2 weeks |
| Anti-Tyrosine Hydroxylase (rabbit, polyclonal) | Merck Millipore (AB152) | 3705678 | AB_390204 | 1:400 → 100 | 2 weeks |
| Recombinant Alexa Fluor® 647 Anti-GFAP (rabbit, monoclonal) | Abcam (ab194325) | GR342668-1 | NA | 1:2000 → 250 | 4 weeks |
| Recombinant Anti-Cytokeratin 7 (rabbit, monoclonal) | Abcam (ab181598) | 1015350-30 | AB_2783822 | 1:2000 → 500 | 2 weeks |
| Recombinant Anti-Cytokeratin 19 (rabbit, monoclonal) | Abcam (ab52625) | GR3384962-5 | AB_2281020 | 1:800 → 200 | 2 weeks |
| Alexa Fluor® 647 AffiniPure Bovine Anti-Goat Secondary | Jackson Immuno Research (805-605-180) | 156773 | AB_2340885 | 1:400 → 100 | 2 weeks |
| Alexa Fluor® 647 AffiniPure Donkey Anti-Mouse Secondary | Jackson Immuno Research (715-605-150) | 155989 | AB_2340862 | 1:1000 → 250,<br>1:800 → 200,<br>1:200 → 50 | 2 weeks |
| **Alexa Fluor™ 647 Donkey anti-Rabbit Highly Cross-Adsorbed Secondary** | Invitrogen (A-31573) | 2359136 | AB_2536183 | 1:200 → 50 | 2 weeks |
| Alexa Fluor® 647 AffiniPure Donkey Anti-Sheep Secondary | Jackson Immuno Research (713-605-147) | 2335729 | AB_2340751 | 1:800 → 200 | 2 weeks |
| Cy3™ AffiniPure Donkey Anti-Goat Secondary | Jackson Immuno Research (115-165-003) | 149607 | AB_2340813 | 1:800 → 200 | 2 weeks |
| Cy3™ AffiniPure Donkey Anti-Rabbit Secondary | Jackson Immuno Research (711-165-152) | 155995 | AB_2307443 | 1:2000 → 500,<br>1:1000 → 250,<br>1:800 → 200,<br>1:400 → 100 | 2 weeks |

### Supplementary Movies

**Movie S1. Dopaminergic neurons in archival human midbrain.** 3D rendering of an archival human midbrain sample, stained for dopaminergic neurons via anti-tyrosine hydroxylase antibody and imaged at 1x magnification using mesoSPIM V5. Zoom into the substantia nigra and switch to high-magnification imaging (12.6x; mesoSPIM V5) of a region-of-interest (ROI). Navigating through the ROI 3D rendering along the long neurites, branching into a dense fiber network.

**Movie S2. Neuronal tracing in archival human substantia nigra.** 3D rendering of the substantia nigra from an archival human midbrain sample (region-of-interest as shown in Movie S1 and Fig. 3), stained for dopaminergic neurons via anti-tyrosine hydroxylase antibody and imaged at 12.6x magnification using mesoSPIM V5. The movie presents consecutive excerpts of raw mesoSPIM images (xy-planes) before increasing transparency to highlight the three-dimensional structure of the dendrites. Three traced dopaminergic neurons with their main branches are shown using color coding.

**Movie S3. Cortical neurons in archival human brain.** 3D rendering of an archival human brain cortex sample, stained for neurons via anti-NeuN antibody and imaged at 1.25x magnification using mesoSPIM V5. Zoom into the cortex and switch to high-magnification imaging (12.6x; mesoSPIM V5) of a region-of-interest (ROI), highlighting layers I-IV. Driving through the ROI 3D stack slice by slice along the z-axis, illustrating the large number of human cortical neurons in this sample, with distinct architecture of pyramidal and granular neurons.

**Movie S4. Microglia and vessels in archival human hippocampus.** 3D rendering of an archival human hippocampus sample, stained for microglia (cyan) via anti-IBA1 antibody and vessels (magenta) via anti-podocalyxin antibody and imaged at 0.8x magnification using mesoSPIM V5. Zoom into the middle of the sample and switch to high-magnification imaging (12.6x; mesoSPIM V5) of a region-of-interest (ROI). Navigating through the ROI 3D rendering, highlighting microglia morphology with their short processes close or attached to blood vessels. Zoom-out and switch back to 3D rendering at 0.8x magnification, going back to the start position.

**Movie S5. Nerve fibers in archival human skin.** 3D rendering of an archival human skin sample, stained for nerve fibers via anti-PGP9.5 antibody and imaged at 1.6x magnification using mesoSPIM V5. Zoom into the dermis and navigating along the long sensory nerve fibers up to a Pacinian corpuscle. Zoom-out and navigating to the epidermis. Zoom-in, turn, and switch to 4x magnification imaging (mesoSPIM V5) of a region-of-interest (ROI) in the transversal view. Navigating through the ROI 3D rendering up to the thin sensory nerve fibers, which innervate the epidermis, and zoom-out.

**Movie S6. Glomeruli and tubules in archival human kidney.** 3D rendering of an archival human kidney sample, stained for glomeruli (blue) via anti-nephrin antibody and for tubules (cyan) via anti-E-cadherin antibody, and imaged at 1x magnification using mesoSPIM V5. Zoom into the middle of the sample up to 3x magnification. Driving through the 3D stack slice by slice along the z-axis, illustrating the large number of glomeruli in this kidney sample. Further zoom into one glomerulus and switch to high-magnification imaging (12.6x; mesoSPIM V5), highlighting glomerulus morphology with Bowman capsule, outgoing proximal tubule, and surrounding distal tubule by driving through the region-of-interest (ROI) 3D stack slice by slice along the z-axis. Switch to 3D rendering of the ROI and zoom-out. Switch back to 3D rendering at 1x magnification and further zoom-out up to the starting position.

**Movie S7. Detected cortical neurons in archival human brain.** Stitched and 4x-downsampled 3D stack of an archival human brain cortex sample, stained for neurons via anti-NeuN antibody and imaged at 4x magnification using mesoSPIM V5. Driving through the 3D stack slice by slice along the z-axis, while zooming into a region-of-interest (ROI), highlighting cortical layers III and IV with pyramidal and granular neurons, respectively. Overlay of the raw 3D stack with detected neurons (yellow) and driving through the 3D stack slice by slice along the z-axis to illustrate the overlap between originally stained and detected neurons.

## References

1. N. Sukswai, J. D. Khoury, Immunohistochemistry Innovations for Diagnosis and Tissue-Based Biomarker Detection. Curr Hematol Malig Rep 14, 368–375 (2019).

2. M. Ittmann, Anatomy and Histology of the Human and Murine Prostate. Cold Spring Harb Perspect Med 8, (2018).

3. F. Blum, Über Formaldehyd als Härtungsmittel. Zentralblatt für allgemeine Pathologie und pathologische Anatomie 4, 1–14 (1893).

4. J. Sy, L. C. Ang, Microtomy: Cutting Formalin-Fixed, Paraffin-Embedded Sections. Methods Mol Biol 1897, 269–278 (2019).

5. K. Tainaka, A. Kuno, S. I. Kubota, T. Murakami, H. R. Ueda, Chemical Principles in Tissue Clearing and Staining Protocols for Whole-Body Cell Profiling. Annu Rev Cell Dev Biol 32, 713–741 (2016).

6. H. R. Ueda et al., Tissue clearing and its applications in neuroscience. Nat Rev Neurosci 21, 61–79 (2020).

7. A. N. Davison, M. Wajda, Analysis of lipids from fresh and preserved adult human brains. Biochem J 82, 113–117 (1962).

8. M. Inoue, R. Saito, A. Kakita, K. Tainaka, Rapid chemical clearing of white matter in the post-mortem human brain by 1,2-hexanediol delipidation. Bioorg Med Chem Lett 29, 1886–1890 (2019).

9. L. M. Molina et al., LiverClear: A versatile protocol for mouse liver tissue clearing. STAR Protoc 3, 101178 (2022).

10. N. Renier et al., iDISCO: a simple, rapid method to immunolabel large tissue samples for volume imaging. Cell 159, 896–910 (2014).

11. M. Belle et al., Tridimensional Visualization and Analysis of Early Human Development. Cell 169, 161–173.e112 (2017).

12. C. R. Scalia et al., Antigen Masking During Fixation and Embedding, Dissected. J Histochem Cytochem 65, 5–20 (2017).

13. T. Liebmann et al., Three-Dimensional Study of Alzheimer’s Disease Hallmarks Using the iDISCO Clearing Method. Cell Rep 16, 1138–1152 (2016).

14. N. Tanaka et al., Whole-tissue biopsy phenotyping of three-dimensional tumours reveals patterns of cancer heterogeneity. Nat Biomed Eng 1, 796–806 (2017).

15. S. Hildebrand, A. Schueth, A. Herrler, R. Galuske, A. Roebroeck, Scalable Labeling for Cytoarchitectonic Characterization of Large Optically Cleared Human Neocortex Samples. Sci Rep 9, 10880 (2019).

16. A. Schueth et al., Efficient 3D light-sheet imaging of very large-scale optically cleared human brain and prostate tissue samples. Commun Biol 6, 170 (2023).

17. S. Zhao et al., Cellular and Molecular Probing of Intact Human Organs. Cell 180, 796–812.e719 (2020).

18. H. Mai et al., Scalable tissue labeling and clearing of intact human organs. Nat Protoc 17, 2188–2215 (2022).

19. H. M. Lai et al., Next generation histology methods for three-dimensional imaging of fresh and archival human brain tissues. Nat Commun 9, 1066 (2018).

20. A. K. L. Liu, H. M. Lai, R. C. Chang, S. M. Gentleman, Free of acrylamide sodium dodecyl sulphate (SDS)-based tissue clearing (FASTClear): a novel protocol of tissue clearing for three-dimensional visualization of human brain tissues. Neuropathol Appl Neurobiol 43, 346–351 (2017).

21. F. Perbellini et al., Free-of-Acrylamide SDS-based Tissue Clearing (FASTClear) for three dimensional visualization of myocardial tissue. Sci Rep 7, 5188 (2017).

22. K. Chung et al., Structural and molecular interrogation of intact biological systems. Nature 497, 332–337 (2013).

23. M. Morawski et al., Developing 3D microscopy with CLARITY on human brain tissue: Towards a tool for informing and validating MRI-based histology. Neuroimage 182, 417–428 (2018).

24. E. Murray et al., Simple, Scalable Proteomic Imaging for High-Dimensional Profiling of Intact Systems. Cell 163, 1500–1514 (2015).

25. Y. G. Park et al., Protection of tissue physicochemical properties using polyfunctional crosslinkers. Nat Biotechnol, (2018).

26. Y. H. Lin et al., Revealing intact neuronal circuitry in centimeter-sized formalin-fixed paraffin-embedded brain. Elife 13, (2024).

27. T. Ku et al., Elasticizing tissues for reversible shape transformation and accelerated molecular labeling. Nat Methods 17, 609–613 (2020).

28. S. Nojima et al., CUBIC pathology: three-dimensional imaging for pathological diagnosis. Sci Rep 7, 9269 (2017).

29. F. F. Voigt et al., The mesoSPIM initiative: open-source light-sheet microscopes for imaging cleared tissue. Nat Methods 16, 1105–1108 (2019).

30. N. Vladimirov et al., Benchtop mesoSPIM: a next-generation open-source light-sheet microscope for cleared samples. Nat Commun 15, 2679 (2024).

31. Y. Tan, C. P. L. Chiam, Y. Zhang, H. L. Tey, L. G. Ng, Research Techniques Made Simple: Optical Clearing and Three-Dimensional Volumetric Imaging of Skin Biopsies. J Invest Dermatol 140, 1305–1314.e1301 (2020).

32. S. A. Schnell, W. A. Staines, M. W. Wessendorf, Reduction of lipofuscin-like autofluorescence in fluorescently labeled tissue. J Histochem Cytochem 47, 719–730 (1999).

33. E. A. Susaki et al., Advanced CUBIC protocols for whole-brain and whole-body clearing and imaging. Nat Protoc 10, 1709–1727 (2015).

34. H. Hama et al., ScaleS: an optical clearing palette for biological imaging. Nat Neurosci 18, 1518–1529 (2015).

35. R. Cai et al., Panoptic imaging of transparent mice reveals whole-body neuronal projections and skull-meninges connections. Nat Neurosci 22, 317–327 (2019).

36. F. Preusser et al., FRC-QE: a robust and comparable 3D microscopy image quality metric for cleared organoids. Bioinformatics 37, 3088–3090 (2021).

37. K. M. Boergens et al., webKnossos: efficient online 3D data annotation for connectomics. Nat Methods 14, 691–694 (2017).

38. R. C. M. Vulders, R. C. van Hoogenhuizen, E. van der Giessen, P. J. van der Zaag, Clearing-induced tisssue shrinkage: A novel observation of a thickness size effect. PLoS One 16, e0261417 (2021).

39. Z. Molnár et al., New insights into the development of the human cerebral cortex. J Anat 235, 432–451 (2019).

40. S. R. y. Cajal, Comparative study of the sensory areas of the human cortex. (Clark University, 1899).

41. G. Leuba, L. J. Garey, Evolution of neuronal numerical density in the developing and aging human visual cortex. Hum Neurobiol 6, 11–18 (1987).

42. R. Gittins, P. J. Harrison, Neuronal density, size and shape in the human anterior cingulate cortex: a comparison of Nissl and NeuN staining. Brain Res Bull 63, 155–160 (2004).

43. S. Berg et al., ilastik: interactive machine learning for (bio)image analysis. Nat Methods 16, 1226–1232 (2019).

44. N. Renier et al., Mapping of Brain Activity by Automated Volume Analysis of Immediate Early Genes. Cell 165, 1789–1802 (2016).

45. J. Kabat, P. Król, Focal cortical dysplasia - review. Pol J Radiol 77, 35–43 (2012).

46. ILAE Commission Report. The epidemiology of the epilepsies: future directions. International League Against Epilepsy. Epilepsia 38, 614–618 (1997).

47. T. Bast, G. Ramantani, A. Seitz, D. Rating, Focal cortical dysplasia: prevalence, clinical presentation and epilepsy in children and adults. Acta Neurol Scand 113, 72–81 (2006).

48. I. Blümcke et al., The clinicopathologic spectrum of focal cortical dysplasias: a consensus classification proposed by an ad hoc Task Force of the ILAE Diagnostic Methods Commission. Epilepsia 52, 158–174 (2011).

49. I. Blumcke et al., Neocortical development and epilepsy: insights from focal cortical dysplasia and brain tumours. Lancet Neurol 20, 943–955 (2021).

50. S. Chong, J. H. Phi, J. Y. Lee, S. K. Kim, Surgical Treatment of Lesional Mesial Temporal Lobe Epilepsy. J Epilepsy Res 8, 6–11 (2018).

51. P. Widdess-Walsh, B. Diehl, I. Najm, Neuroimaging of focal cortical dysplasia. J Neuroimaging 16, 185–196 (2006).

52. M. Heers et al., Spectral bandwidth of interictal fast epileptic activity characterizes the seizure onset zone. Neuroimage Clin 17, 865–872 (2018).

53. A. K. Glaser et al., A hybrid open-top light-sheet microscope for versatile multi-scale imaging of cleared tissues. Nat Methods 19, 613–619 (2022).

54. Y. Tan et al., 3-Dimensional Optical Clearing and Imaging of Pruritic Atopic Dermatitis and Psoriasis Skin Reveals Downregulation of Epidermal Innervation. J Invest Dermatol 139, 1201–1204 (2019).

55. S. Abadie et al., 3D imaging of cleared human skin biopsies using light-sheet microscopy: A new way to visualize in-depth skin structure. Skin Res Technol 24, 294–303 (2018).

56. J. Jeong et al., An optimized visualization and quantitative protocol for in-depth evaluation of lymphatic vessel architecture in the liver. Am J Physiol Gastrointest Liver Physiol 325, G379–G390 (2023).

57. C. C. Chen et al., Human liver afferent and efferent nerves revealed by 3-D/Airyscan super-resolution imaging. Am J Physiol Endocrinol Metab 326, E107–E123 (2024).

58. T. Yoshizawa et al., Three-dimensional analysis of ductular reactions and their correlation with liver regeneration and fibrosis. Virchows Arch 484, 753–763 (2024).

59. W. B. Fabyan et al., LiverMap pipeline for 3D imaging of human liver reveals volumetric spatial dysregulation of cirrhotic vasculobiliary architecture. bioRxiv, (2024).

60. C. Adori et al., Disorganization and degeneration of liver sympathetic innervations in nonalcoholic fatty liver disease revealed by 3D imaging. Sci Adv 7, (2021).

61. C. Kong et al., Multiscale and Multimodal Optical Imaging of the Ultrastructure of Human Liver Biopsies. Front Physiol 12, 637136 (2021).

62. K. R. Weiss, F. F. Voigt, D. P. Shepherd, J. Huisken, Tutorial: practical considerations for tissue clearing and imaging. Nat Protoc 16, 2732–2748 (2021).

63. M. Thom et al., Cortical neuronal densities and lamination in focal cortical dysplasia. Acta Neuropathol 110, 383–392 (2005).

64. I. C. Galvão et al., Identifying cellular markers of focal cortical dysplasia type II with cell-type deconvolution and single-cell signatures. Sci Rep 13, 13321 (2023).

65. J. Schindelin et al., Fiji: an open-source platform for biological-image analysis. Nat Methods 9, 676–682 (2012).

66. B. Schmid, J. Schindelin, A. Cardona, M. Longair, M. Heisenberg, A high-level 3D visualization API for Java and ImageJ. BMC Bioinformatics 11, 274 (2010).

67. B. Schmid, P. Tripal, Z. Winter, R. Palmisano, 3Dscript.server: true server-side 3D animation of microscopy images using a natural language-based syntax. Bioinformatics 37, 4901–4902 (2021).

68. R. C. Team. (R Foundation for Statistical Computing, Vienna, Austria., 2020).

69. A. Bria, G. Iannello, TeraStitcher - a tool for fast automatic 3D-stitching of teravoxel-sized microscopy images. BMC Bioinformatics 13, 316 (2012).

70. G. Van Rossum, & Drake Jr, F. L. (Centrum voor Wiskunde en Informatica Amsterdam, 1995).

71. G. Van Rossum, & Drake, F. L. (Scotts Valley, CA: CreateSpace, 2009).

72. A. L. Tyson et al., A deep learning algorithm for 3D cell detection in whole mouse brain image datasets. PLoS Comput Biol 17, e1009074 (2021).

73. A. e. a. Paszke. (Advances in Neural Information Processing Systems, 2019), vol. 32, pp. 8024–8035.

74. J. Duchi, Hazan, E., & Singer, Y. (Journal of machine learning research, 2011), vol. 12.

75. S. Fort, Huiyi Hu, and Balaji Lakshminarayanan. (arXiv preprint, 2019), vol. arXiv:1912.02757.

76. P. A. Yushkevich et al., User-guided 3D active contour segmentation of anatomical structures: significantly improved efficiency and reliability. Neuroimage 31, 1116–1128 (2006).

